# MAnorm2 for quantitatively comparing groups of ChIP-seq samples

**DOI:** 10.1101/2020.01.07.896894

**Authors:** Shiqi Tu, Mushan Li, Fengxiang Tan, Haojie Chen, Jian Xu, David J. Waxman, Yijing Zhang, Zhen Shao

**Affiliations:** CAS Key Laboratory of Computational Biology, CAS-MPG Partner Institute for Computational Biology, Shanghai Institute of Nutrition and Health, Shanghai Institutes for Biological Sciences, Chinese Academy of Sciences, Shanghai 200031, China; University of Chinese Academy of Sciences, Beijing 100049, China; Children’s Medical Center Research Institute, Department of Pediatrics, University of Texas Southwestern Medical Center, Dallas, TX 75390, USA; Department of Biology and Bioinformatics Program, Boston University, Boston MA 02215, USA; National Key Laboratory of Plant Molecular Genetics, CAS Center for Excellence in Molecular Plant Sciences, Shanghai Institute of Plant Physiology and Ecology, Shanghai Institutes for Biological Sciences, Chinese Academy of Sciences, Shanghai 200032, China

**Author notes:** **Corresponding authors:** Dr. Zhen Shao.

## Abstract

Eukaryotic gene transcription is regulated by a large cohort of chromatin associated proteins, and inferring their differential binding sites between cellular contexts requires a rigorous comparison of the corresponding ChIP-seq data. We present MAnorm2, a new computational tool for quantitatively comparing groups of ChIP-seq samples. MAnorm2 uses a hierarchical strategy to normalize ChIP-seq data and then performs differential analysis by assessing within-group variability of ChIP-seq signals under an empirical Bayes framework. In this framework, MAnorm2 considers the abundance of differential ChIP-seq signals between groups of samples and the possibility of different within-group variability between groups. When samples in each group are biological replicates, MAnorm2 can reliably identify differential binding events even between highly similar cellular contexts. Using a number of real ChIP-seq data sets, we observed that MAnorm2 clearly outperformed existing tools for differential ChIP-seq analysis, with the improvement in performance being most dramatic when the groups of samples being compared had distinct global within-group variability.

## Introduction

Chromatin immunoprecipitation followed by high-throughput sequencing (ChIP-seq) has become the premier technology for profiling the genome-wide localization of chromatin-binding proteins, including transcription factors and histones with various modifications [1, 2]. An important downstream analysis of ChIP-seq data is to identify the genomic regions that are associated with differential binding intensities between different cellular contexts, which is essential to understanding the underlying mechanisms orchestrating the dynamics of gene expression program during various biological processes, including development and the onset of disease [3–5]. With the decrease of sequencing costs, researchers now incline to perform differential ChIP-seq analysis between groups of samples. On one hand, when the samples of each group are biological replicates for the same ChIP-seq experiment, differential analysis on group level can achieve much better specificity and sensitivity than between individual samples [6]. This improvement is especially vital for comparing ChIP-seq samples from heterogeneous tissues or closely related cell lineages [7–9]. On the other hand, when analyzing ChIP-seq profiles for tissues or cells obtained from different individuals, researchers may choose to classify them according to the age, sex, health status or disease subtype of each donor, and then perform differential analysis between groups of profiles to identify differential binding events associated with the group characteristics. This analysis is of particular interest in the studies of personal epigenomes, where fluctuations of histone modification levels across humans are often of functional importance and are best understood on population level [10–13].

Despite the importance of differential ChIP-seq analysis on group level, it remains a highly challenging computational task to reliably assess on a genome-wide scale the statistical significances of observed differences in ChIP-seq signal intensities between groups of samples, owing to the high level of noise and variability intrinsic to ChIP-seq data [6, 14]. In general, the success of a group-level differential ChIP-seq analysis relies on a robust approach for normalizing multiple ChIP-seq samples, as well as a sophisticated statistical model for accurately assessing the variability of ChIP-seq signals across samples of the same group (referred to as within-group variability) [15, 16]. Here we present a new computational tool for differential ChIP-seq analysis between groups of samples. The new tool, named MAnorm2, has made specific efforts to tackle the above two challenges in a manner that accounts for ChIP-seq data specific characteristics.

Regarding normalization, in practice, signal-to-noise (S/N) ratio can vary significantly across different ChIP-seq samples [6, 16], which greatly increases the difficulty of normalization. We previously developed MAnorm for normalizing a pair of ChIP-seq samples. It alleviates the problem of S/N ratio by using common peaks (genomic regions enriched with ChIP-seq reads) of the two samples to infer a reference model for globally normalizing ChIP-seq signals [17]. This strategy has also been exploited by later methods for differential ChIP-seq analysis [18]. In MAnorm2, we extended MAnorm to normalization of any number of ChIP-seq samples. We further incorporated a hierarchical strategy into the normalization of groups of samples to account for the similarity structure among samples. Specifically, by first normalizing samples separately within each group and then performing a between-group normalization, MAnorm2 could improve both the unbiasedness and robustness.

As for assessing within-group variability, in the field of differential RNA-seq analysis, the strategy of borrowing information between genes with similar expression levels under an empirical Bayes framework has been adopted by several cutting-edge methods, including limma-trend, voom and DESeq2 [19–21]. Among them, limma-trend fits a mean-variance curve (MVC) and squeezes gene-wise variance estimates towards the curve; voom is similar to limma-trend except that it encodes the fitted MVC into the precision weights of different measurements; DESeq2 uses the negative binomial distribution to model read counts and aims at fitting a mean-dispersion curve (MDC). In practice, these methods borrow strength between genes to improve variance/dispersion estimation, which could compensate for the lack of sufficient replicates. The same principle applies to ChIP-seq data as well, and many studies have directly used these methods to perform differential ChIP-seq analysis [13, 22–24]. Despite the usefulness of modeling mean-variance/dispersion trend under an empirical Bayes framework, no methods exploiting this strategy have been specifically developed for ChIP-seq data. As ChIP-seq data are typically associated with much higher noise level and variability than RNA-seq data, the statistical models originally designed for differential RNA-seq analysis may not be flexible enough to account for the characteristics of ChIP-seq data.

One problem is that the above RNA-seq methods derive the mean gene expression levels for fitting an MVC/MDC by taking the average across all individual samples, disregarding their group labels. This strategy generally applies well to differential RNA-seq analysis, since in most cases the vast majority of genes are expected to have non-differential expression between biological conditions. For differential ChIP-seq analysis, however, the strategy may considerably bias the resulting MVC/MDC owing to the abundance of differential ChIP-seq signals. In practice, differential protein-binding events can be abundant even between very similar cellular contexts [10, 25]. One of the reasons is that the activity of regulatory elements, especially those at distal regions, is generally more variable across cellular contexts than is gene expression [10, 26, 27]. Another problem is that the RNA-seq methods derive a single variance/dispersion estimate for each gene, without explicitly modeling the difference in within-group variability between different groups. In practice, however, within-group variability of ChIP-seq signals could vary significantly across groups. For instance, when comparing ChIP-seq profiles between normal individuals and cancer patients, the within-group variability in the cancer group is often much higher than that in the normal group. In MAnorm2, we have made specific efforts to address these concerns. In particular, we resolve the two problems by deriving mean and variance estimates separately within each group of samples, adjusting the variance estimates from different groups based on the global within-group variability of each group, and pooling the resulting mean-variance pairs into a regression process.

We compared the performance of MAnorm2, limma-trend, voom and DESeq2 on a number of real ChIP-seq data sets. For each differential ChIP-seq analysis that we conducted, MAnorm2 performed as well or better than the other three methods, with the improvement in performance being most dramatic when the two groups of samples being compared had distinct global within-group variability. We also made a comparison between MAnorm2 and two existing methods (ChIPComp [18] and PePr [28]) for differential ChIP-seq analysis, and found they were clearly outperformed by MAnorm2.

## Results

### Hierarchical normalization for groups of ChIP-seq samples

To facilitate the understanding of the working principle of MAnorm2, we first give a brief description of some of its basic concepts. An MAnorm2 analysis starts with a count matrix and an occupancy matrix. The rows and columns of both matrices correspond to a pre-defined list of genomic intervals and a set of ChIP-seq samples, respectively. For each genomic interval in each sample, the count matrix records the number of sequencing reads mapped to the interval, and the occupancy matrix uses a binary variable to indicate whether the interval is enriched with reads (i.e., whether it is a peak region). Formally, we refer to a genomic interval as occupied by a ChIP-seq sample if the interval is enriched with reads in the sample.

For each MAnorm2 analysis performed in this study, we compiled the list of genomic intervals by separately calling peaks for each related ChIP-seq sample, merging all the resulting peaks, and dividing up broad merged peaks into consecutive genomic bins (narrow ones were left untouched). We then determined the occupancy status of each resulting genomic interval for each sample based on the overlap between the interval and the peaks identified for the sample. See Methods for details about the construction of input matrices of MAnorm2, but note that the MAnorm2 machinery is independent of the exact way of coming up with a list of genomic intervals and defining their occupancy states.

We previously developed MAnorm for pairwise comparison of ChIP-seq samples. It normalizes two individual samples by removing the overall M-A trend (M and A values refer to log2 fold change and average log2 read count, respectively) observed at their common peaks, based on the assumption of no global changes of protein-binding intensities at these regions [17]. MAnorm has been shown to give a better performance in handling ChIP-seq samples with distinct S/N ratios than the methods originally designed for normalizing microarray and RNA-seq data [16, 17]. In MAnorm2, we retained the core assumption of MAnorm, and further devised a hierarchical scheme for the normalization of groups of samples to take advantage of the similarity structure among samples.

Here we use the normalization of H3K4me3 ChIP-seq data for two human lymphoblastoid cell lines (LCLs) as an example. The two LCLs, referred to as GM12891 and GM12892, were derived from different Caucasian individuals, and each cell line was associated with three biological replicates (data obtained from Kasowski et al. [10]). In the hierarchical normalization process, MAnorm2 first separately normalizes the replicates for each LCL and then applies a between-group normalization (Fig. 1A). For the first step, MAnorm2 selects one sample from each group as its baseline (based on the size factors of samples by default; see Methods), and repeatedly normalizes every other sample of the group against that baseline. Specifically, to normalize each non-baseline sample against the corresponding baseline sample, MAnorm2 applies a linear transformation to the log2 read counts of the non-baseline sample, such that the M-A trend observed at the common peak regions of the two samples is removed (see Methods; note also that MAnorm2 considers the genomic intervals that are occupied by both samples to be their common peak regions). For the second step, MAnorm2 creates a reference ChIP-seq profile for each LCL by taking the average across all of its replicates (using normalized log2 read counts determined in the first step), and re-applies the above procedure for within-group normalization to the resulting two reference profiles. Then, the linear transformation derived for the non-baseline reference profile is equally applied to each individual replicate of the LCL it represents. The only technical problem of this approach is that we need to determine the occupancy states of genomic intervals for each reference profile (or equivalently, each group of samples). In this study, we considered a genomic interval to be occupied by a group of samples if it was occupied by at least one sample in the group.

**Figure 1.**
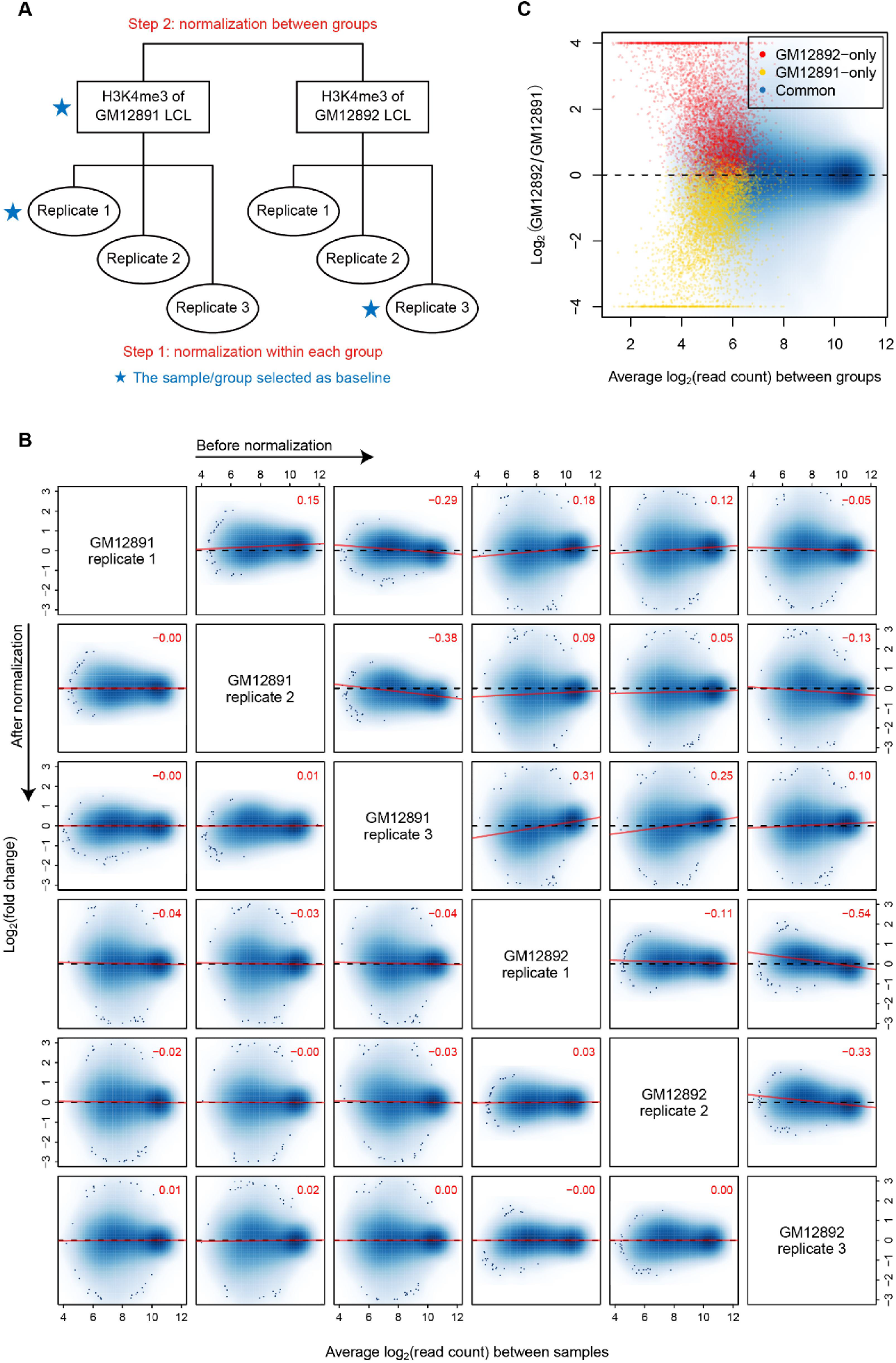
Hierarchical normalization for groups of ChIP-seq samples. **(A)** Diagram illustrating the hierarchical normalization process applied to the H3K4me3 ChIP-seq samples of GM12891 and GM12892 lymphoblastoid cell lines (LCLs). **(B)** MA scatter plots for each pair of the H3K4me3 samples before (upper triangular part) and after (lower triangular part) normalization. Only common peak regions of the associated two samples are used to draw each plot, and the numeric value in the top-right corner gives the Pearson correlation coefficient (PCC) between M and A values across these regions. Solid red lines are fitted by using the least squares estimation. **(C)** MA scatter plot for the normalized reference H3K4me3 profiles of GM12891 and GM12892 LCLs. Here the genomic intervals are classified based on their occupancy states in the two groups of samples.

After completing the entire normalization process, we found that the M-A trend at the common peak regions of each pair of samples was largely eliminated, independent of whether the two samples came from the same group or whether they had ever been selected as baseline (Fig. 1B). This finding indicated that the normalized ChIP-seq signal intensities were comparable across all six samples. And we could then quantify the fold changes of H3K4me3 levels between the two LCLs by calculating M values between their (normalized) reference profiles. Of note, the M values for LCL-specific peak regions were systematically biased towards the corresponding LCL (Fig. 1C), which contrasted with the roughly symmetric distribution of the M values for common peak regions and supported the need to remove sample/group specific peak regions from “globally invariant” regions when performing a normalization.

### Modeling mean-variance trend under an empirical Bayes framework

After normalization, MAnorm2 models the normalized log2 read counts as following the normal distribution. This strategy is similar to the one employed by voom, except that voom first normalizes read counts (based on the total read count of each sample) and then applies log2 transformation [20]. For a differential analysis between two groups of samples, a simple and straightforward method would be to apply a *t*-test to the normalized signal intensities of each genomic interval. In practice, however, the number of ChIP-seq samples in each group is usually small (two or three for the data sets used in this study), which may compromise the statistical power of *t*-tests for identifying differential binding events, considering the large uncertainty of the variance estimate for each interval that is derived from a very limited number of signal measurements.

To tackle this problem, MAnorm2 borrows strength between genomic intervals with similar signal levels. Specifically, it calculates observed means and variances for individual intervals within each group of samples, adjusts the observed variances from different groups to make them comparable across groups, and pools the resulting mean-variance pairs into a regression process to fit an MVC (Fig. 2A; see also Methods and Supplementary Note 1). The fitted MVC is then incorporated into the differential analysis under an empirical Bayes framework for achieving shrinkage for variance estimation. Similar to limma-trend [19, 20], MAnorm2 specifies an inverse-gamma distribution as the prior distribution of the variance of each genomic interval, with the associated parameters determined by the MVC and a hyper-parameter denoted by *d*_0_, which is referred to as the number of prior degrees of freedom and amounts to the number of additional samples acquired by sharing information between intervals. In effect, the final variance estimate for each individual interval is a weighted average among the prior variance (obtained from the MVC) and the (adjusted) observed variances from different groups, with the weights being proportional to their respective numbers of degrees of freedom (see Methods).

**Figure 2.**
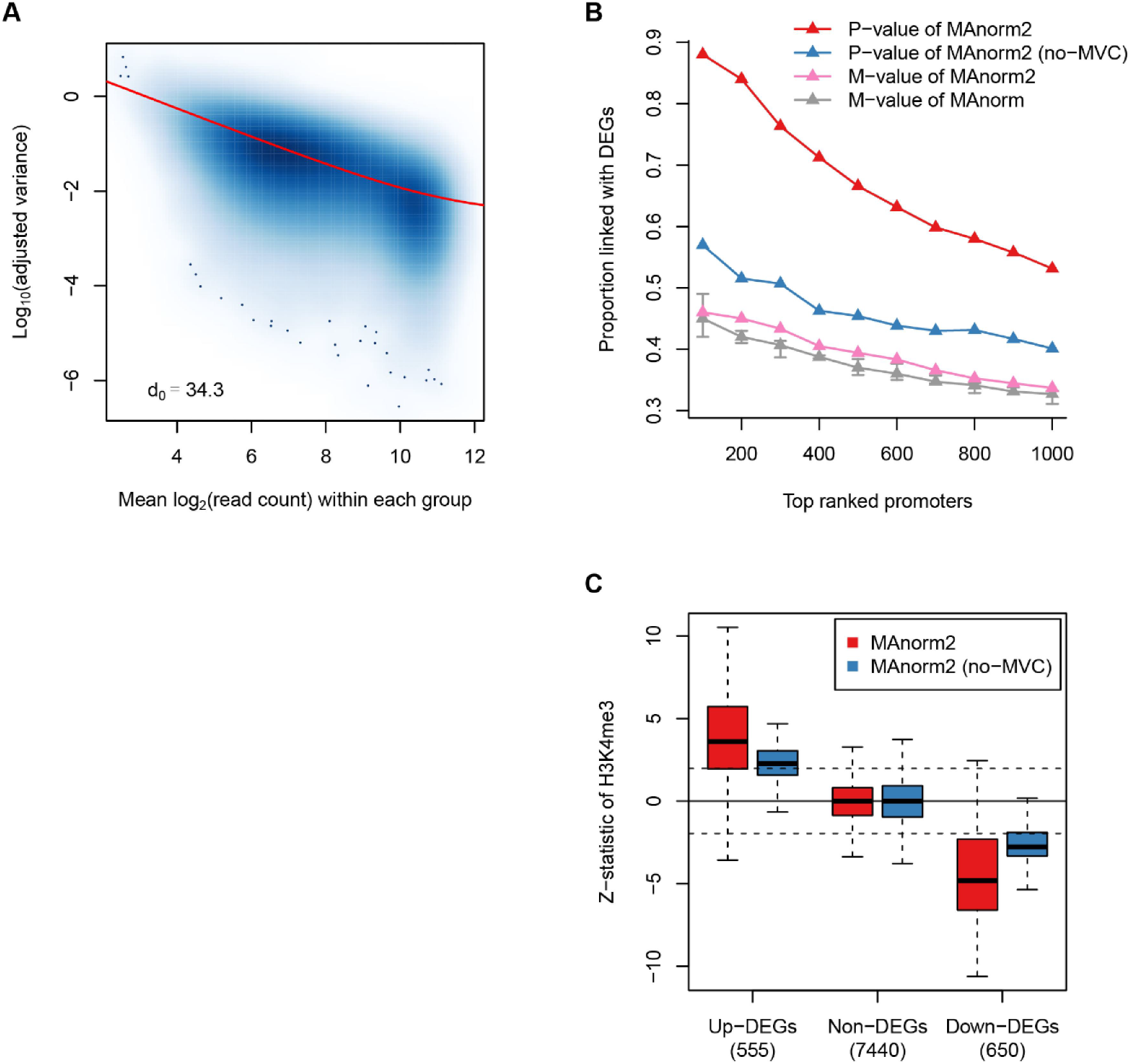
Modeling mean-variance trend to improve variance estimation. **(A)** Scatter plot showing the mean-variance trend associated with the H3K4me3 ChIP-seq samples of GM12891 and GM12892 LCLs. Red line depicts the fitted mean-variance curve (MVC), and *d*_0_ gives the corresponding number of prior degrees of freedom. **(B)** For the identification of differential H3K4me3 levels between GM12891 and GM12892 LCLs, the proportion of true discoveries among top ranked genomic intervals at gene promoters is plotted against the number of top ranked intervals. Here true discoveries are defined as the intervals that are linked with differentially expressed genes (DEGs), which are identified by applying DESeq2 to the corresponding RNA-seq data with a *p*-value cutoff of 0.01. MAnorm has been applied to all possible comparisons between an individual H3K4me3 ChIP-seq sample of GM12891 and an individual sample of GM12892, and we show here the median as well as the first and third quartiles of the true discovery proportions achieved across the total nine comparisons. **(C)** Box plots for the *z*-statistic equivalents of the *p*-values assigned to the genomic intervals located at the promoter regions of DEGs and non-DEGs. Dotted lines correspond to a two-tailed *p*-value of 0.05. Here non-DEGs are defined as those genes having a DESeq2 *p*-value larger than 0.5 and a fold change less than 2.

By conducting a comparison of the H3K4me3 ChIP-seq samples between GM12891 and GM12892 LCLs, we evaluated the performance of several MAnorm/MAnorm2 statistics for calling differential genomic intervals. These statistics included the M values derived by MAnorm and MAnorm2 as well as the *p*-values derived by MAnorm2 and one of its variants, referred to as no-MVC. Technically, no-MVC directly uses observed variances as the final variance estimates and is intrinsically equivalent to a *t*-test. Since H3K4me3 levels at promoter regions of genes are strongly correlated with their transcription levels, we defined, for the genomic intervals located at gene promoters, true differential ones as those that were linked with differentially expressed genes (DEGs), which were identified by applying DESeq2 [21] to the corresponding RNA-seq data. Using each of the statistics, we ranked the promoter intervals in order of evidence of having differential H3K4me3 levels between the two LCLs, and calculated the proportions of true discoveries among top ranked intervals (Fig. 2B). In this way, we found that, compared with the M values from MAnorm, those from MAnorm2 were associated with an overall increase in the DEG proportions, suggesting that generating biological replicates can lead to more reliable estimates of fold changes of ChIP-seq signals. Regarding *p*-values, from no-MVC to MAnorm2, we observed a substantial improvement in accurately picking out the true differential intervals, which illustrated the importance of using sophisticated statistical techniques to improve variance estimation. We also examined the exact *p*-values assigned to the genomic intervals located at the promoter regions of DEGs and non-DEGs (Fig. 2C). For better graphical presentation, shown in Figure 2C are actually *z*-statistic equivalents of the *p*-values, which were obtained by mapping the *p*-values together with the signs of M values to the standard normal distribution [29]. For example, a *z*-statistic of 1.96 corresponds to a two-tailed *p*-value about 0.05. It can be seen that, benefiting from the modeling of mean-variance relationship for reducing the uncertainty of variance estimates, MAnorm2 considerably increased the sensitivity for identifying differential intervals, and it maintained the specificity without arbitrarily biasing the *p*-values of the promoters of non-DEGs towards significance.

The strategy of shrinking variance/dispersion estimates under an empirical Bayes framework has been adopted by several existing tools for differential RNA-seq analysis [20, 21]. Technically, these tools improve the adaptivity to data sets of various characteristics by empirically determining the shrinkage strength. Similarly, our MAnorm2 uses the hyper-parameter *d*_0_ to effectively control the degree to which observed variances are squeezed towards MVC, and *d*_0_ is estimated from the data set under analysis. Despite the wide use of this strategy, few studies have used concrete examples to demonstrate specifically how it contributes to the adaptivity of a method. In particular, the advantage of empirical Bayes shrinkage over using directly prior variances/dispersions is of much interest. Here, we applied MAnorm2 to differential ChIP-seq analysis in two scenarios where the associated variance structures were of different complexity, by incorporating H3K4me3 ChIP-seq data for two additional Caucasian LCLs (referred to as GM12890 and SNYDER, each associated with two biological replicates [10]). In the first scenario, we compared the H3K4me3 samples of GM12890 with those of SNYDER. In the second scenario, we classified the total four LCLs into male and female ones to perform a between-sex comparison. Note that, for this analysis, we used a reference H3K4me3 profile to represent each LCL and conducted a two-versus-two comparison as in the first scenario. The variance structure in the second scenario is clearly more complicated than that in the first scenario, as the within-group variability in the second scenario is additionally associated with the epigenetic variation across human individuals [10]. This difference is also indicated by the distinct *d*_0_ estimates derived by MAnorm2 in the two analyses (*d*_0_ estimates are 14.6 and 4.2 in the first and the second scenario, respectively). We then used the corresponding gene expression data to assess the performance of MAnorm2 and another variant of it, which is referred to as MVC-only and uses directly prior variances as the final variance estimates. In summary, while the performance of MVC-only is comparable with that of MAnorm2 in the first scenario, it is significantly outperformed by MAnorm2 in the second one (Supplementary Fig. 1). These results suggest that MVC-only is only suited to data sets with a highly regular variance structure. They also explicitly demonstrate how empirical Bayes shrinkage for variance estimation improves the adaptivity of a method. See Supplementary Note 2 for a much more detailed discussion of this topic.

### Comparing MAnorm2 with other empirical Bayes methods that account for mean-variance/dispersion relationship

As mentioned above, the strategy of modeling mean-variance/dispersion trend under an empirical Bayes framework has been adopted by several methods that are initially developed for differential RNA-seq analysis (e.g., limma-trend, voom and DESeq2 [19–21]), and many studies have applied them to differential ChIP-seq analysis by first processing ChIP-seq data into a regular count matrix [13, 22–24]. The primary differences between MAnorm2 and these RNA-seq methods relate to two considerations of MAnorm2: (i) MAnorm2 calculates observed means and variances of signal intensities separately within each group of samples to make the unbiasedness of MVC fitting resistant to the abundance of differential signals between groups, and (ii) MAnorm2 adjusts the observed variances from different groups of samples to make them comparable across groups. Technically, for the latter, MAnorm2 introduces *γ_j_*, referred to as a variance ratio factor, to parameterize the global within-group variability of group *j*. When comparing samples between group 1 and 2, MAnorm2 derives an estimate of *γ*_2_/*γ*_1_ and uses the estimated ratio to adjust the observed variances from group 2 (see Methods and Supplementary Note 1).

To investigate how MAnorm2 benefits from the above two considerations, we made a systematic comparison of MAnorm2, limma-trend, voom and DESeq2 in differential analysis of practical ChIP-seq data. We first compared the performance of all the four methods in identifying genomic intervals with differential H3K4me3 levels between GM12891 and GM12892 LCLs. This analysis can serve as a good example to demonstrate alone the effect of the first consideration, as there is only a small difference in global within-group variability between the corresponding two groups of ChIP-seq samples (the estimated *γ_GM_*_12892_/*γ_GM_*_12891_ from MAnorm2 is 0.802). This observation can be explained by the fact that the ChIP-seq samples of each group are biological replicates for an individual LCL and all the involved samples are generated from the same study. Based on the corresponding gene expression data, we found that, compared with the other three methods, MAnorm2 provided a better ranking of the genomic intervals that are located at gene promoters (Fig. 3A).

**Figure 3.**
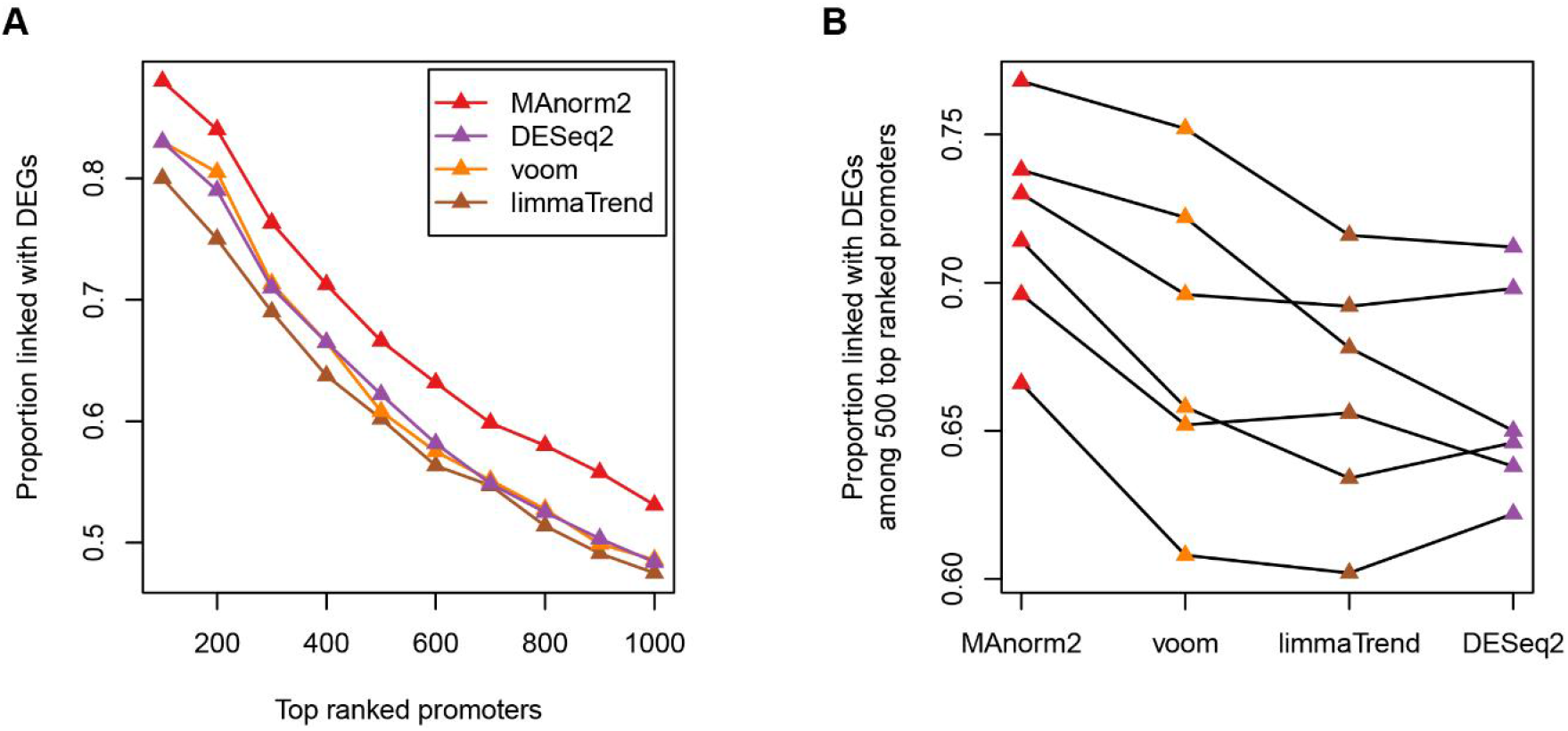
Comparison among empirical Bayes methods that take mean-variance/dispersion relationship into account. **(A)** MAnorm2 as well as three existing empirical Bayes methods that model mean-variance/dispersion relationship has been applied to the identification of differential H3K4me3 levels between GM12891 and GM12892 LCLs. For each of the methods, the proportion of true discoveries among top ranked genomic intervals at gene promoters is plotted against the number of top ranked intervals. **(B)** We performed all possible pairwise comparisons of H3K4me3 levels among GM12890, GM12891, GM12892 and SNYDER LCLs, and we show here the true discovery proportions among 500 top ranked intervals (at gene promoters) achieved by each method. In the plot, each line corresponds to an individual comparison between two LCLs, and the methods are sorted by the average true discovery proportion across the six comparisons.

We further performed all possible pairwise comparisons of H3K4me3 ChIP-seq data among the four LCLs (GM12890, GM12891, GM12892 and SNYDER). For each of the resulting six comparisons, we selected 100, 200 up to 1000 top ranked genomic intervals at promoter regions and examined the proportions of true discoveries among them. Figure 3B gives the true discovery proportions among 500 top ranked intervals achieved by each method, and it can be seen that MAnorm2 outperforms the other three methods in all the comparisons. More detailedly, each method is associated with 60 proportions of true discoveries, and MAnorm2 provides the best performance in 51 of the cases (including 2 cases where MAnorm2 ties with voom; Supplementary Fig. 2). These results support the necessity for taking the abundance of differential ChIP-seq signals into account when fitting an MVC.

To investigate the practical utility of the second consideration of MAnorm2, we performed a differential analysis of H3K27ac ChIP-seq data between normal human individuals and cancer patients. In practice, H3K27ac ChIP-seq profiles for cancer patients are typically associated with much higher variability than those for normal individuals, owing to the heterogeneity of cancer tissues/cells as well as the diversity of subtypes and stages of the disease. We first collected H3K27ac ChIP-seq data for three human chronic lymphocytic leukemia (CLL) cell lines (referred to as MEC1, OSU-CLL and CII) that are derived from different patients [13]. For the normal counterparts, we collected data for GM12891, GM12892 and SNYDER LCLs [10], which were selected to match the sex composition of the CLL group. Note that all the LCLs and CLL cell lines are generated by using Epstein-Barr virus to transform primary B cells harvested from donors and, thus, the comparison between the LCL group and the CLL group makes clear biological sense.

Using MAnorm2, we normalized the H3K27ac ChIP-seq signals across the six cell lines. Principle component analysis (PCA) on the resulting normalized signal intensities showed that the global within-group variability of the CLL group is indeed much higher than the LCL group (Fig. 4A). We have also drawn a scatter plot that maps the observed means and variances (of normalized signal intensities) from each group (Fig. 4B). Overall, data points from the CLL group have a clear tendency to lie above those from the LCL group, indicating an increased magnitude of signal variances associated with the CLL group. Consistently, MAnorm2 derived an estimate of *γ_CLL_*/*γ_LCL_* about 3.66, indicating that the global within-group variability of the CLL group is over three-fold compared to the LCL group. The estimated ratio was then used by MAnorm2 to adjust the observed variances from the CLL group, and an MVC was subsequently fitted on the resulting mean-variance pairs from both groups (Fig. 4C). The whole process is essentially to normalize observed variances from different groups, and MAnorm2 manages to integrate this normalization into a statistical model for the following differential analysis (see Methods). Again, based on the corresponding gene expression profiles, we defined true differential genomic intervals among the ones that were located at gene promoters, and we compared the rankings of promoter intervals provided by different methods (Fig. 4D). We found that MAnorm2 clearly outperformed the other three methods, which indicated the importance of accounting for the difference in global within-group variability between different groups of samples when performing a group-level differential ChIP-seq analysis.

**Figure 4.**
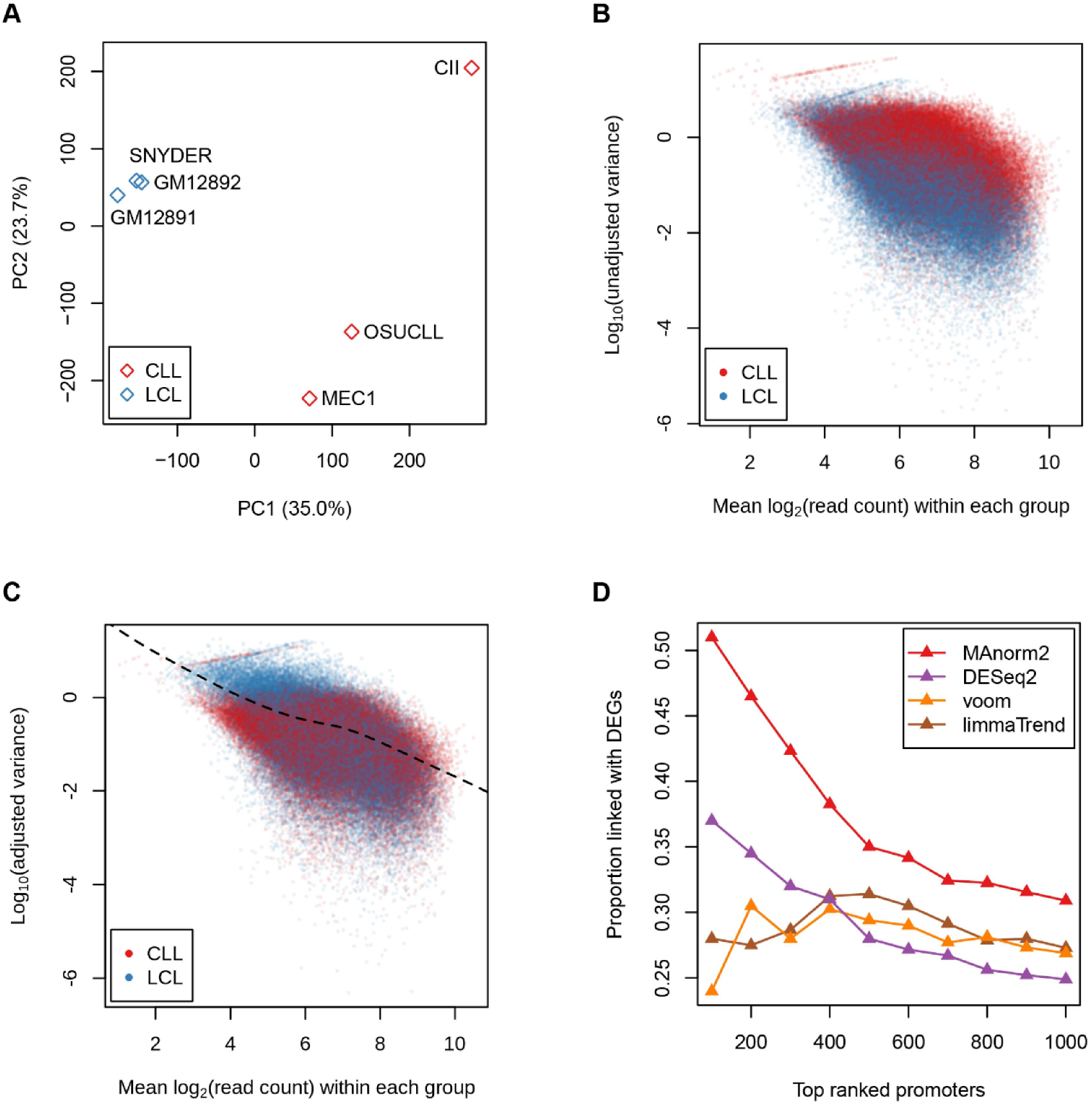
Explicitly modeling the difference in global within-group variability between different groups. **(A)** Principle component analysis (PCA) was applied to the normalized H3K27ac ChIP-seq signal intensities (derived by MAnorm2) of three LCLs and three chronic lymphocytic leukemia (CLL) cell lines. Shown here is a scatter plot for the first two principle components. **(B)** Scatter plot of log10 observed variances against observed mean signal intensities of individual genomic intervals. Here the variances and means are calculated separately from each group of cell lines. **(C)** Scatter plot of log10 adjusted variances against observed mean signal intensities. **(D)** MAnorm2 and three other methods have been used to call differential H3K27ac ChIP-seq signals between the two groups of cell lines. For each method, the proportion of true discoveries among top ranked genomic intervals at gene promoters is plotted against the number of top ranked intervals.

### Comparing MAnorm2 with other tools for group-level differential ChIP-seq analysis

We selected two representative computational tools for group-level differential ChIP-seq analysis to compare with MAnorm2. The two tools, named ChIPComp [18] and PePr [28], represent two broad classes of methods for differential ChIP-seq analysis [6, 14]. More specifically, ChIPComp requires its users to provide pre-defined peaks for each single ChIP-seq sample while PePr has no such requirement (we could see that MAnorm2 and ChIPComp are actually of the same class of methods with respect to the dependence on pre-defined peaks). For each of the differential analyses of H3K4me3 ChIP-seq data between a pair of LCLs and the differential analysis of H3K27ac ChIP-seq data between LCLs and CLL cell lines, we compared the ranking of genomic intervals at gene promoters provided by MAnorm2 with those from ChIPComp and PePr. Note that we have separately used each of two statistics derived by ChIPComp to rank intervals, which were *p*-value and the posterior probability for a fold change of ChIP-seq signal to be larger than 2 [18]. We observed that MAnorm2 gives the highest proportion of true discoveries in all cases (Fig. 5 and Supplementary Fig. 3). Together, these comparisons between MAnorm2 and the tools for differential ChIP/RNA-seq analysis demonstrate the power of modeling mean-variance trend under an empirical Bayes framework and indicate the usefulness of adapting this modeling strategy to the characteristics of ChIP-seq data.

**Figure 5.**
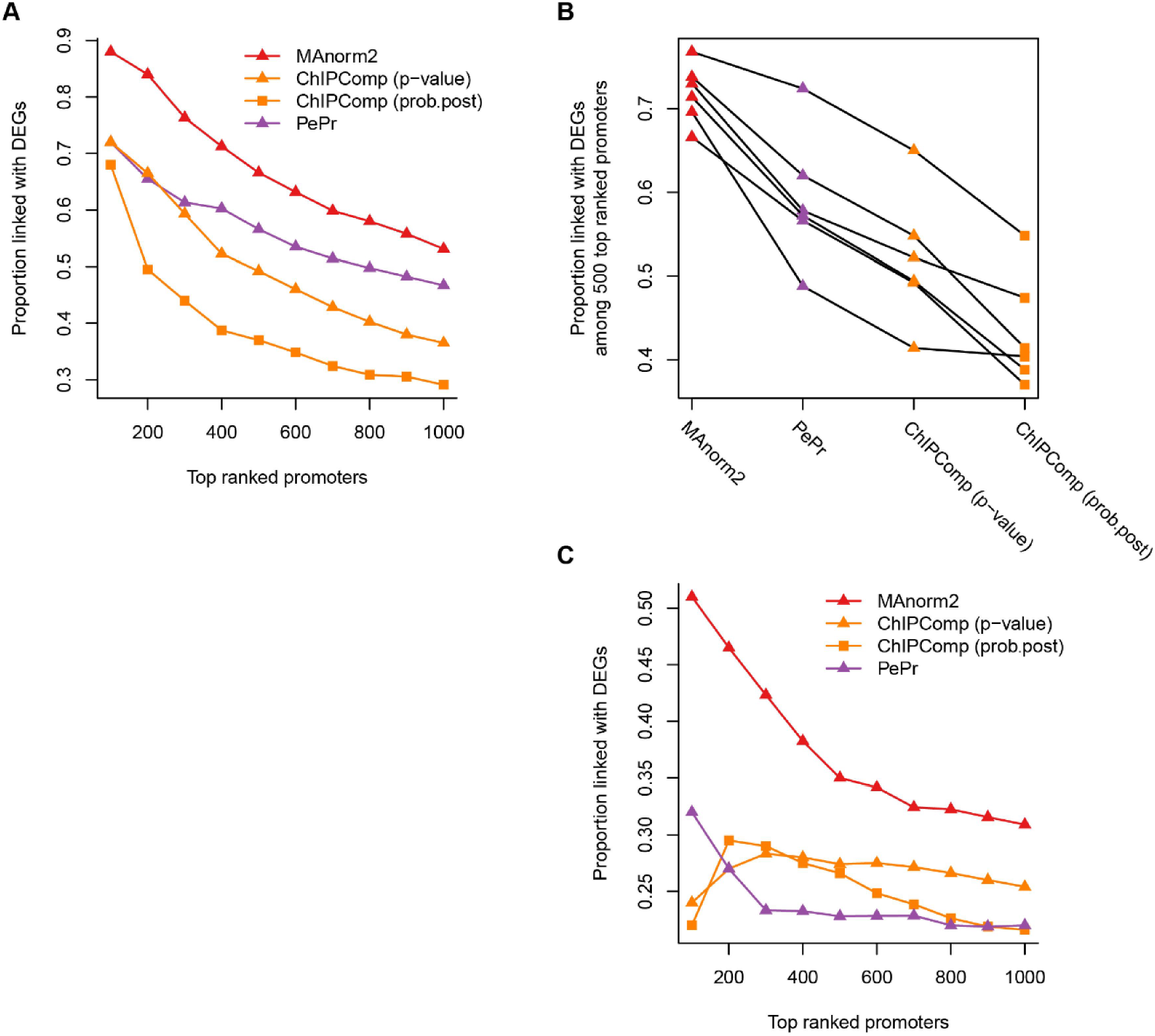
Comparing MAnorm2 with other tools for differential ChIP-seq analysis on group level. **(A)** Comparison of the rankings of genomic intervals at gene promoters provided by different methods in the differential analysis of H3K4me3 ChIP-seq data between GM12891 and GM12892 LCLs. **(B)** Comparison of the proportions of true discoveries among 500 top ranked promoter intervals achieved by different methods in each of the differential analyses of H3K4me3 ChIP-seq data between a pair of LCLs. **(C)** Comparison of the rankings of promoter intervals from different methods in the differential analysis of H3K27ac ChIP-seq data between LCLs and CLL cell lines.

## Discussion

In the study, we developed MAnorm2 for quantitatively comparing groups of ChIP-seq samples. Technically, MAnorm2 is comprised of a hierarchical normalization scheme and an empirical Bayes model that exploits mean-variance relationship to improve variance estimates. We have demonstrated the standard workflow of each of the two parts for comparing two groups of samples, but both parts could be extended in several ways for practical ChIP-seq data analysis.

For the normalization part, the hierarchical scheme is designed for taking advantage of the similarity structure among samples. It could also increase the stability of normalization results by avoiding selecting a single baseline sample from many samples. In the study, we have shown how to apply a hierarchical normalization to two groups of samples corresponding to different cellular contexts. In theory, the normalization method could be generalized to samples sorted into an arbitrary hierarchical structure. More specifically, by traversing the associated hierarchical tree from bottom to top, we could iteratively normalize the samples associated with each tree branch and create a reference profile to represent the branch. Similar to the two-group normalization, this normalization process always chooses the samples/profiles most resembling each other to normalize at each round of iteration. For practical data analysis, we expect that increasing the complexity of hierarchical tree to account for additional factors affecting the similarity structure among samples (e.g., batch effects) may help reducing normalization biases, especially for large-scale studies where a great number of samples are involved [11, 12].

As for the statistical model part, MAnorm2 uses a multivariate normal (MVN) distribution to model the normalized signal intensities of each genomic interval in each group of samples (in the study, normalized signal intensities derived from the normalization part of MAnorm2 were all normalized log2 read counts). Technically, for each interval in each group, the covariance matrix of the MVN distribution is expressed as a symmetric matrix (termed structure matrix) times a scalar that quantifies the within-group variability (see Methods). Although all the structure matrices used in this study were simply identity matrices, the users could use structure matrix to easily incorporate existing tools for modeling the precision weights of signal measurements from different samples as well as the correlations among the measurements [20, 29], which, for example, may help dealing with ChIP-seq samples that are associated with distinct quality and/or batch effects. Another extension of the MAnorm2 model regards comparison of more than two groups of samples. Under the MVN framework, the model can be readily extended to simultaneous comparison of any number of groups (Supplementary Note 3). Here we use the differential analysis of H3K4me3 ChIP-seq data among GM12890, GM12891, GM12892 and SNYDER LCLs as example. For comparison, we also applied one-way analysis of variance (ANOVA) to the normalized signal intensities and calculated maximum fold changes across the four groups of samples. By identifying DEGs among the four LCLs, we found that MAnorm2 provided a much better ranking of the genomic intervals at gene promoters than the other two methods (Fig. 6A). In addition, compared with ANOVA, MAnorm2 considerably increased the sensitivity for identifying differential intervals without sacrificing the specificity (Fig. 6B).

**Figure 6.**
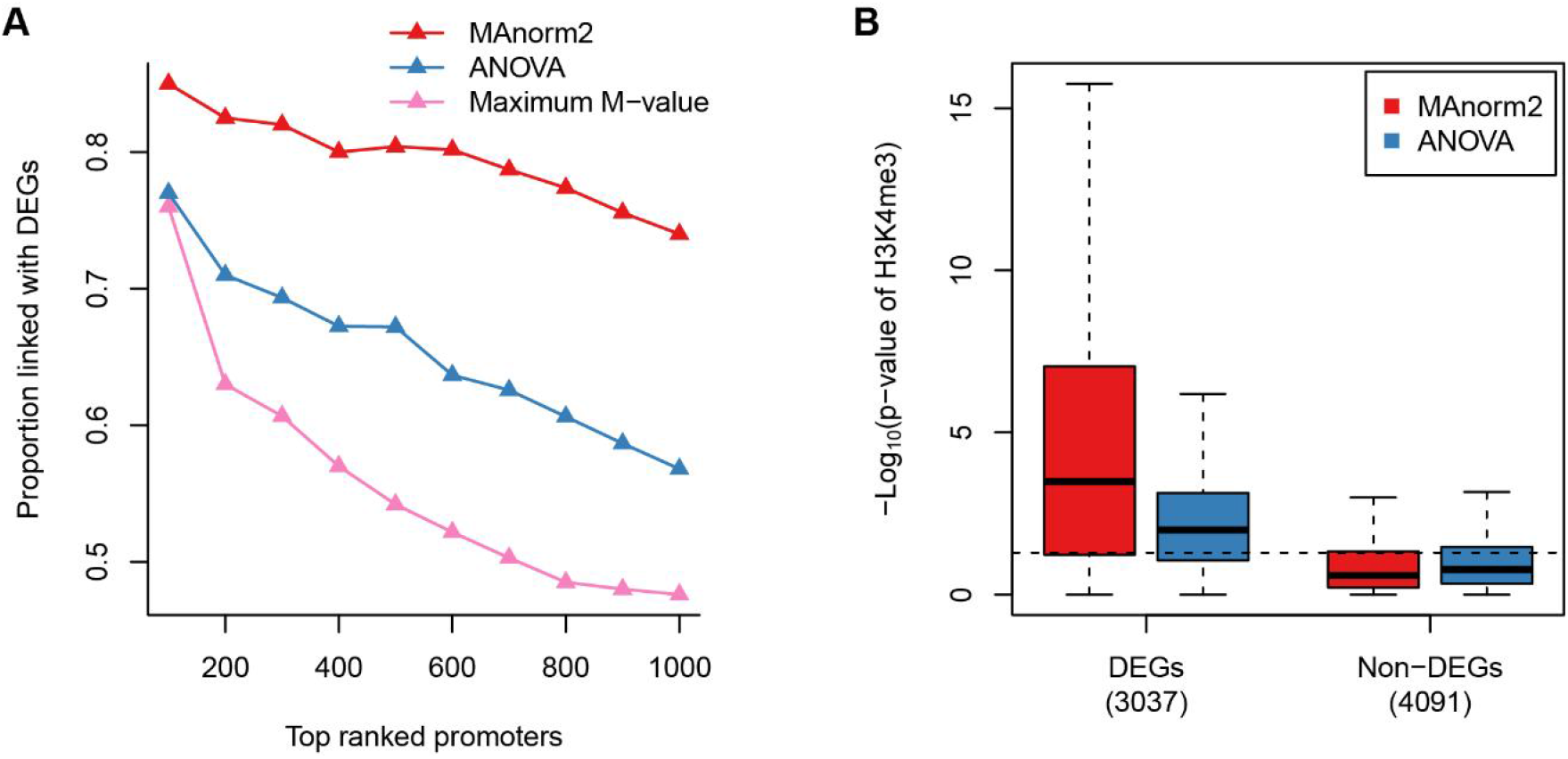
Simultaneously comparing multiple groups of ChIP-seq samples. **(A)** For the identification of differential H3K4me3 levels among GM12890, GM12891, GM12892 and SNYDER LCLs, the proportion of true discoveries among top ranked genomic intervals at gene promoters is plotted against the number of top ranked intervals. Here DEGs are identified by applying the likelihood ratio test provided by DESeq2 to the corresponding four groups of RNA-seq samples, with a *p*-value cutoff of 0.01. **(B)** Box plots for the –log10 *p*-values assigned to the genomic intervals located at the promoter regions of DEGs and non-DEGs. Dotted line corresponds to a *p*-value of 0.05. Here non-DEGs are defined as those genes having a DESeq2 *p*-value larger than 0.5 and a maximum fold change across the four groups less than 2.

The two parts of MAnorm2 are relatively independent of each other, and they both operate on continuous variables (thus, the input signal measurements of MAnorm2 are not limited to integers). Together, these properties make it easy to incorporate existing methods for adjusting for confounding factors into an MAnorm2 analysis. For example, MAnorm2 has not devised a specific method for correcting for background signals that are measured by input samples. In practice, the users could perform such a correction before invoking the normalization part of MAnorm2, by separately subtracting raw/normalized input read counts from each ChIP-seq sample [22, 28] or by modeling the relationship between background and ChIP-seq signals in a more sophisticated manner [18]. Another example relates to batch effects. Most existing tools for removing batch effects are initially developed for continuous measurements from microarray experiments [30, 31]. If the ChIP-seq data under analysis are found to be associated with serious batch effects, one could apply these tools to the normalized signal intensities from MAnorm2 before invoking its modeling part for the following differential analysis.

## Supporting information

Supplementary Software

## Acknowledgements

We thank Dr. Guo-Cheng Yuan and Dr. Stuart H. Orkin in Dana Farber Cancer Institute for helpful suggestions. This work was supported by the National Basic Research Program of China (2018YFA0107602), the National Natural Science Foundation of China (31871280 and 31701140), the “100-Talent Program” of Chinese Academy of Science (Y516C11851 to Z.S.) and the U.S. National Institutes of Health (NIH grant DK121998 to D.J.W.).

## Author Contributions

S.T. and Z.S. conceived the study; S.T. and M.L. developed the algorithms; S.T. analyzed the data with help from M.L. and F.T.; Z.S. supervised the study; S.T. and Z.S. wrote the manuscript with contributions from all the other authors.

## Methods

### Data sets and preprocessing

H3K4me3 ChIP-seq data and RNA-seq data for four human lymphoblastoid cell lines (LCLs; GM12890, GM12891, GM12892 and SNYDER) derived from different Caucasian individuals were obtained from Kasowski et al. [10]. H3K27ac ChIP-seq data for GM12891, GM12892 and SNYDER LCLs were obtained from the same study. H3K27ac ChIP-seq data and RNA-seq data for three human chronic lymphocytic leukemia (CLL) cell lines (MEC1, OSU-CLL and CII) were obtained from Ott et al. [13].

We started processing the ChIP-seq and RNA-seq samples with raw sequencing reads. We first applied quality and adapter trimming to the 3’ end of every raw read by using Trim Galore [32]. The resulting ChIP-seq and RNA-seq reads were then aligned to the hg19 reference genome by bowtie [33] and STAR [34], respectively. To avoid potential artefacts from PCR amplification [35], we separately removed duplicates for each individual sample, which means we kept at most one read (for each single-end sequencing sample) or read pair (for each paired-end sequencing sample) at each genomic location. Note that the genomic location of each single read was considered to be the coordinate and strand of its 5’ end in the reference genome, and the genomic location of each read pair was considered to be the coordinates of the associated two 5’ ends in plus and minus strands respectively.

### Processing read alignments and calling peaks for ChIP-seq samples

We separately identified peaks for each of the ChIP-seq samples by using MACS 1.4 [36]. The ‘--shiftsize’ parameter of MACS 1.4 controls the distance in bp by which the genomic coordinate of the 5’ end of each aligned read is shifted downstream to reach the exact protein binding site, which is presumed to be the center of the underlying DNA fragment from which the read is sequenced. When using MACS 1.4, the single value specified for ‘--shiftsize’ is universally applied to all reads, but the sizes of collected DNA fragments for sequencing do vary. To make full use of the parameter, for each aligned read pair of a paired-end sequencing sample, we inferred the center of the underlying DNA fragment as the midpoint between the coordinates of the associated two 5’ ends, and the read pair was then converted into a single read whose 5’ end lies upstream of the inferred center with an exact distance of 100 bp. We also noticed that the setting of ‘--shiftsize’ could influence the resulting peak lengths. To unify the general peak length across different ChIP-seq samples, we have processed single-end sequencing samples in a similar manner. Specifically, for each aligned read of a single-end sequencing sample, the associated DNA fragment center was inferred as the site lying downstream of the read’s 5’ end with a certain distance, which was half the fragment size used for size-selecting the library (this information was available for all the single-end ChIP-seq samples used in the study), and the read was then shifted such that the resulting 5’ end lies exactly 100 bp upstream of the inferred center. Finally, for each processed ChIP-seq sample, peak calling was performed against the corresponding input sample with the parameters ‘-g hs --nomodel --shiftsize=100 --keep-dup=all’ (we had already removed duplicates for each of the ChIP-seq samples before applying the above processing). Note that input samples were available for all the ChIP-seq samples used in the study and they were processed in exactly the same way as the ChIP-seq samples. To be clear, all the differential ChIP-seq analyses presented in the study were based on the same processed read alignments as used for peak calling.

### Input matrices of MAnorm2

For performing a differential analysis between two or more groups of ChIP-seq samples, MAnorm2 takes a count matrix and an occupancy matrix as input. The rows of both matrices correspond to a pre-defined list of genomic intervals, and their columns correspond to the involved ChIP-seq samples. For each genomic interval in each sample, the count matrix records the number of sequencing reads that fall within the interval, and the occupancy matrix uses a binary variable to indicate whether the interval is enriched with reads (or whether the interval is a peak region). Note that MAnorm2 refers to a genomic interval as occupied by a ChIP-seq sample if the interval is enriched with reads in the sample.

We have specifically developed a software package named MAnorm2_utils for integrating ChIP-seq data into a regular table suited as input of MAnorm2. For each MAnorm2 analysis performed in the study, we compiled the two input matrices by applying MAnorm2_utils 1.0.0 to the (processed) read alignment results of associated ChIP-seq samples and the peaks identified separately for each of them, with the parameters ‘--typical-bin-size=2000 --shiftsize=100 --keep-dup=all --filter=blacklist.bed’. Here we give a brief description of how MAnorm2_utils works under the parameter settings (see https://github.com/tushiqi/MAnorm2_utils/tree/master/docs for full documentation of MAnorm2_utils). First, it merges all the provided peaks of different samples and deduces a summit for each merged peak. It then divides up each merged peak into consecutive non-overlapping genomic bins of 2 kb, by aligning the center of the first bin with the deduced summit and extending the bin list towards both directions until the merged peak is completely covered. For each edge bin, if its center is covered by the merged peak, it is retained and trimmed to the corresponding peak edge. Otherwise, it is trimmed and absorbed into its predecessor. We have chosen 2 kb as the bin size in this study, which was for matching the typical peak length observed in the H3K4me3 and H3K27ac ChIP-seq samples. The resulting genomic intervals are then filtered to remove the ones that overlap the blacklisted regions [37]. At this moment, MAnorm2_utils has finished deriving the list of genomic intervals corresponding to the rows of both input matrices (in effect, each narrow merged peak, if not overlapping the blacklist, is left untouched and directly serves as a final genomic interval). MAnorm2_utils next shifts downstream the 5’ end of each aligned read by 100 bp to reach the inferred protein binding site, and the resulting sites from each sample are then assigned to the genomic intervals, which determines the count matrix. For the occupancy matrix, MAnorm2_utils considers a genomic interval to be occupied by a ChIP-seq sample only if the interval’s center is covered by some peak of the sample.

### Normalizing individual ChIP-seq samples

MAnorm2 has designed two routines for normalizing a set of any number of ChIP-seq samples. One of them is for normalizing individual samples. The other is for normalizing multiple groups of samples each having been separately normalized.

To normalize a group of individual ChIP-seq samples, MAnorm2 selects one of them as baseline and normalizes each of the other samples against it. MAnorm2 provides an option for the users to specify the baseline sample by themselves. By default, it uses the median-ratio strategy [38] to derive the size factor of each sample and selects the sample whose log2 size factor is closest to 0 as the baseline (in the study, we always used the default setting). To avoid biases as much as possible, only the genomic intervals that are occupied by all the samples are used to derive their size factors. We next detail the procedure for normalizing a ChIP-seq sample against another.

Suppose that *X* and *Y* are two vectors of log2 read counts (we used an offset of 0.5 in the study) representing the raw signal intensities of two ChIP-seq samples and that we want to normalize *Y* against *X*. Let *x_i_* and *y_i_* be their *i*-th elements, which correspond to the *i*-th genomic interval. MAnorm2 assumes the ChIP-seq signals at the common peak regions of the two samples are globally invariant between them, and it accomplishes the normalization by applying a linear transformation to *Y*. Formally, let 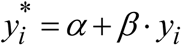 be the normalized signal intensity of interval *i*, where *α* and *β* are coefficients to be determined. We define normalized M and A values as 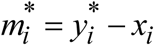 and 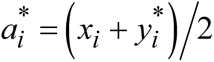, respectively, and we determine the two coefficients by imposing the following two constraints:

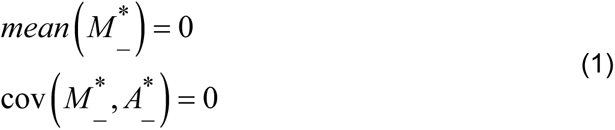

where *mean* and cov refer to sample mean and sample covariance, respectively, and _ indicates the quantities are calculated with respect to the common peak regions (i.e., the genomic intervals occupied by both samples). The solutions for *α* and *β* are given by

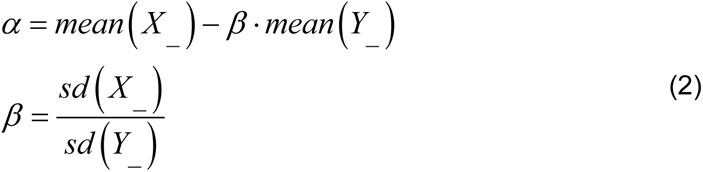

where *sd* refers to sample standard deviation.

### Modeling and normalizing groups of ChIP-seq samples

MAnorm2 models a group of ChIP-seq samples that have already been normalized by using the multivariate normal (MVN) distribution. Suppose *X* is an *n* × *m* matrix recording the normalized signal intensities (i.e., normalized log2 read counts) at *n* genomic intervals for *m* ChIP-seq samples belonging to the same biological condition. Let *X_i_* be a column vector that represents the transpose of the *i*-th row of *X*. We assume

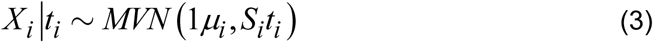

Here *μ_i_* and *t_i_* are two unknown scalars that parametrize the mean signal intensity of interval *i* in this biological condition and the associated signal variation level, respectively; 1 is a column vector of ones; *S_i_*, termed structure matrix, is an *m* × *m* matrix designed for the convenience of incorporating existing tools for modeling the precision weights of signal measurements from different samples as well as the correlations among the measurements [20, 29, 39]. All structure matrices used in the study were simply identity matrices. MAnorm2 next derives mean and variance estimates by applying the generalized least squares method:

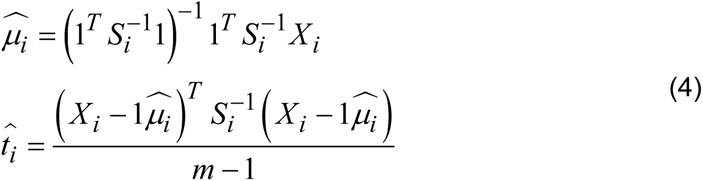

Finally, MAnorm2 uses the vector of 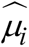 as a reference ChIP-seq profile that represents this group of *m* samples.

For normalizing multiple groups of ChIP-seq samples each having been separately normalized, MAnorm2 first derives a reference profile for each group of samples and determines the occupancy states of genomic intervals in each group. In this study, a genomic interval was considered to be occupied by a group of samples if it was occupied by at least one of the samples. MAnorm2 then applies the procedure for normalizing individual ChIP-seq samples to the resulting reference profiles. Finally, the transformation coefficients derived for each non-baseline reference profile are equally applied to each sample of the corresponding group. Note that, since the transformations are linear, the original structure matrices remain valid for the transformed signals. When comparing two groups of ChIP-seq samples from different biological conditions, MAnorm2 also calculates M values of genomic intervals between their (normalized) reference profiles, which represent the log2 fold changes between the two conditions.

### Modeling mean-variance trend and identifying differential signals between two groups of ChIP-seq samples

MAnorm2 improves variance estimates for individual genomic intervals by borrowing strength between intervals with similar signal levels. Specifically, MAnorm2 captures the underlying mean-variance dependence by fitting a smooth mean-variance curve (MVC).

For *j* = 1, 2, suppose *X_j_* is an *n* × *m_j_* matrix recording the normalized ChIP-seq signal intensities (i.e., normalized log2 read counts) at *n* genomic intervals for *m_j_* samples belonging to condition *j*. Let *X_i_*_, *j*_ be a column vector that represents the transpose of the *i*-th row of *X_j_*. We now make a comparison between condition 1 and 2. Note that equation (3) and (4) are still valid once we add a subscript *j* to each of the associated variables to indicate its group label. We next assume the MVCs of the two groups of samples have the same shape and differ from each other only by a constant factor. Formally, we define 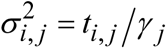, where *γ_j_*, termed variance ratio factor, is a parameter that quantifies the global variation level of ChIP-seq signals across the samples of group *j*. And the complete Bayesian model that takes advantage of mean-variance trend is given by

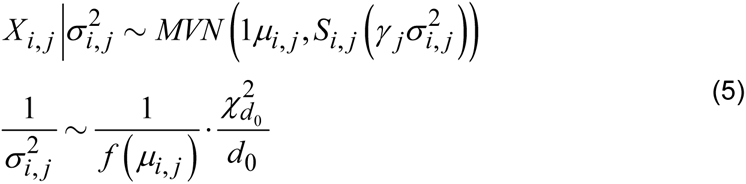

Overall, the above model is similar to limma-trend [19, 20], except that MAnorm2 allows for different global within-group variability between groups of samples. Here *f*(·) refers to the underlying unscaled MVC common to the two groups of samples and *f*(*μ_i,j_*) is called the prior variance of interval *i* in group *j*; *d*_0_ is a hyper-parameter called the number of prior degrees of freedom; 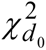 refers to the chi-squared distribution with *d*_0_ degrees of freedom. We also assume that the unscaled variances of non-differential genomic intervals remain invariant across the two groups of samples, which means 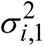 equals 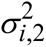 with a probability of one (i.e., they could be treated as the same random variable) for each interval *i* that satisfies *μ_i_*_,1_ = *μ_i_*_,2_. This assumption is consistent with the fact that 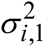 and 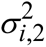 follow the same prior distribution as long as *μ_i_*_,1_ = *μ_i_*_,2_.

Finally, we test the null hypothesis *H*_0_ : *μ_i_*_,1_ = *μ_i_*_,2_ for each genomic interval *i* by using the following key statistic:

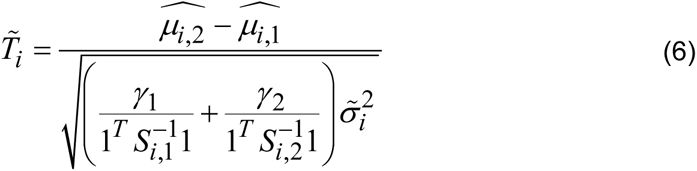

where

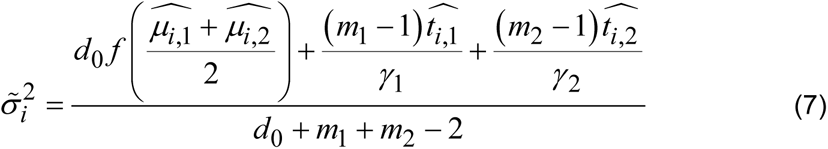

According to the theoretical deduction presented in Smyth et al. [39], if 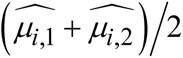 in the above equation were replaced by *μ_i_*_,1_ (or *μ_i_*_,2_), 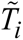 would strictly follow a *t*-distribution under the null hypothesis with (*d*_0_ + *m*_1_ + *m*_2_ − 2) degrees of freedom. Here we derive the mean estimates for determining prior variances by taking the average signal intensities across conditions rather than individual samples, which is for balancing the statistical power for identifying down- and up-regulated signals. Accordingly, a two-sided *p*-value for the statistical test is given by 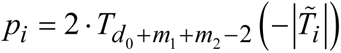, where 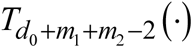 refers to the cumulative distribution function of the *t*-distribution with (*d*_0_ + *m*_1_ + *m*_2_ − 2) degrees of freedom.

As for MVC fitting and parameter estimation, MAnorm2 adopts an empirical Bayes approach in which *f*, *d*_0_, *γ*_1_ and *γ*_2_ are estimated from the data. Supplementary Note 1 gives a fully detailed description of the whole model formulation as well as the associated parameter estimation. Here we stress that, different from many existing empirical Bayes methods that also model mean-variance/dispersion trend [19–21], MAnorm2 fits an MVC by calculating mean and variance estimates separately within each group of samples and adjusting the variance estimates from different groups based on their global within-group variability. More specifically, for fitting *f*, MAnorm2 derives an estimate of *γ*_2_/*γ*_1_ and pools the mean-variance pairs of the form 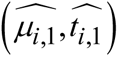 or 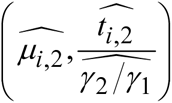 into a weighted gamma-family regression procedure, with ((*m*_1_ − 1) and (*m*_2_ − 1) as the weights of observations from group 1 and 2, respectively.

### Differential analysis of H3K4me3 ChIP-seq data and RNA-seq data between individual LCLs

We performed all possible pairwise comparisons among GM12890, GM12891, GM12892 and SNYDER LCLs. For the differential analysis of H3K4me3 ChIP-seq data, each of the LCLs was associated with two or three biological replicates. For each comparison of H3K4me3 ChIP-seq signals between a pair of LCLs, MAnorm2 was invoked by using MAnorm2_utils to create the input matrices; limma-trend, voom and DESeq2 were applied to the same count matrix as used by MAnorm2; ChIPComp used the associated ChIP-seq samples along with the corresponding input samples and the identified peaks as input. We initialized the core object designed by ChIPComp with the parameters filetype=“bed” and species=“hg19”, and we performed the differential analysis with default settings; PePr used the associated ChIP-seq samples and the corresponding input samples as input, and it was invoked with the parameters ‘-f bed -s 100 -w 1000 --diff --peaktype=sharp -- normalization=inter-group’.

For the differential analysis of RNA-seq data, each of the LCLs was associated with two biological replicates. Aligned reads or read pairs of each RNA-seq sample were assigned to the UCSC annotated genes [40] by htseq-count [41]. To identify differentially expressed genes (DEGs) between each pair of LCLs, we applied DESeq2 to the corresponding count matrix with a *p*-value cutoff of 0.01. For the comparison between GM12891 and GM12892 LCLs, we have also come up with a list of non-DEGs (Fig. 2C), which were defined as the genes that had a DESeq2 *p*-value larger than 0.5 and a fold change (obtained also from DESeq2) less than 2.

### Differential analysis of H3K27ac ChIP-seq data and RNA-seq data between LCLs and CLL cell lines

We performed a comparison between three LCLs (GM12891, GM12892 and SNYDER) and three CLL cell lines (MEC1, OSU-CLL and CII). For the differential analysis of H3K27ac ChIP-seq data, each of the LCLs was associated with two or three biological replicates, and each of the CLL cell lines was associated with a single ChIP-seq sample. For the comparison of H3K27ac levels between the LCL group and the CLL group, MAnorm2_utils was applied to all the associated individual ChIP-seq samples, and MAnorm2 performed the differential analysis by creating a reference H3K27ac profile for each LCL; limma-trend, voom and DESeq2 were invoked by using the count matrix generated by MAnorm2_utils and taking the average read counts (rounded to the nearest integers) across the biological replicates of each LCL; ChIPComp and PePr were invoked by pooling the reads of the biological replicates of each LCL and adopting the same parameter settings as used in the differential analysis of H3K4me3 ChIP-seq data between LCLs (peak calling was re-performed for each pooled sample for invoking ChIPComp).

For the differential analysis of RNA-seq data, each of the LCLs was associated with two biological replicates, and each of the CLL cell lines was associated with three biological replicates. Aligned reads or read pairs of each individual RNA-seq sample were assigned to the UCSC annotated genes by htseq-count. For identifying DEGs between the LCL and CLL groups, we calculated the average read counts (rounded to the nearest integers) across the biological replicates of each cell line and applied DESeq2 to the resulting count matrix with a *p*-value cutoff of 0.01.

### Software availability

We used MAnorm2_utils 1.0.0 and MAnorm2 1.0.0 in the study. MAnorm2_utils 1.0.0 and MAnorm2 1.0.0 are provided as Python and R packages in Supplementary Software, respectively. The latest versions of MAnorm2 and MAnorm2_utils can always be found at https://github.com/tushiqi. MAnorm2_utils has also been uploaded to the PyPI repository (https://pypi.org/project/MAnorm2-utils).

