## Supplementary Notes and Supplementary Figures for "MAnorm2 for quantitatively comparing groups of ChIP-seq samples"

**Contents:**

Supplementary Note 1. Differential analysis between two groups of ChIP-seq samples

- 1.1 Model formulation and hypothesis testing
- 1.2 Mean-variance curve fitting and parameter estimation
- 1.3 Deducing a parametric form for the mean-variance curve

Supplementary Note 2. Being adaptive to the regularity of variance structure

Supplementary Note 3. Simultaneously comparing multiple groups of ChIP-seq samples

References

Supplementary Figure Legends

Supplementary Figures

### Supplementary Note 1. Differential analysis between two groups of ChIP-seq samples

Here we give a detailed description of the statistical model designed in MAnorm2 for calling differential ChIP-seq signals between two groups of samples corresponding to different biological conditions, provided that all the involved samples have been normalized to be comparable with each other.

#### 1.1 Model formulation and hypothesis testing

For $j = 1,2$, suppose $X_j$ is an $n \times m_j$ matrix recording the normalized ChIP-seq signal intensities (i.e., normalized log2 read counts) at $n$ genomic intervals for $m_j$ samples belonging to condition $j$. Let $X_{i,j}$ be a column vector that represents the transpose of row $i$ of $X_j$. We assume

$$X_{i,j} \left| t_{i,j} \sim MVN\left(1\mu_{i,j}, S_{i,j} t_{i,j}\right)\right. \quad (1)$$

Here $MVN$ refers to the multivariate normal distribution; $\mu_{i,j}$ and $t_{i,j}$ are two unknown scalars that parametrize the mean signal intensity of interval $i$ in condition $j$ and the associated signal variation level, respectively; $1$ is a column vector of ones; $S_{i,j}$, termed structure matrix, is an $m_j \times m_j$ matrix designed for the convenience of incorporating existing tools for modeling the precision weights of signal measurements from different samples as well as the correlations among the measurements [1-3]. All structure matrices used in the study were simply identity matrices. MAnorm2 next derives mean and variance estimates by applying the generalized least squares method:

$$\widehat{\mu_{i,j}} = \left(1^T S_{i,j}^{-1} 1\right)^{-1} 1^T S_{i,j}^{-1} X_{i,j}$$
$$\widehat{t_{i,j}} = \frac{\left(X_{i,j} - 1\widehat{\mu_{i,j}}\right)^T S_{i,j}^{-1} \left(X_{i,j} - 1\widehat{\mu_{i,j}}\right)}{m_j - 1} \quad (2)$$

In practice, $m_j$ are typically very small (most ChIP-seq data sets only have two or three biological replicates for each experiment), which results in large uncertainty associated with the $t_{i,j}$ estimators and thus low statistical power for the following differential tests. To improve variance estimates for individual genomic intervals,

MAnorm2 borrows strength between intervals with similar signal levels and captures the underlying mean-variance dependence by fitting a smooth mean-variance curve (MVC). Specifically, it assumes the MVCs of the two groups of samples being compared have the same shape and differ from each other only by a constant factor. Formally, MAnorm2 defines $\sigma_{i,j}^2 = t_{i,j} / \gamma_j$ , where $\gamma_j$ , termed variance ratio factor, is a parameter that quantifies the global variation level of ChIP-seq signals across the samples of group $j$ . Naturally, we have $\widehat{\sigma_{i,j}^2} = \widehat{t_{i,j}} / \gamma_j$ . And the complete Bayesian model that uses mean-variance trend as source of prior information is given by

$$X_{i,j} \left| \sigma_{i,j}^2 \sim MVN\left(1\mu_{i,j}, S_{i,j}\left(\gamma_j \sigma_{i,j}^2\right)\right)\right.$$
$$\frac{1}{\sigma_{i,j}^2} \sim \frac{1}{f\left(\mu_{i,j}\right)} \cdot \frac{\chi_{d_0}^2}{d_0} \quad (3)$$

Overall, the above model is similar to the one described in Smyth et al. [1] as well as its extension [4], except that MAnorm2 allows for different global within-group variability between groups of samples. Specifically, here $f(\cdot)$ refers to the underlying unscaled MVC common to the two groups of samples and $f\left(\mu_{i,j}\right)$ is called the prior variance of interval $i$ in group $j$ ; $\chi_N^2$ refers to the chi-squared distribution with $N$ degrees of freedom; $d_0$, termed the number of prior degrees of freedom, is a hyper-parameter designed for assessing how well in general the variance of an individual interval could be predicted by its mean signal intensity. In practice, $d_0$ amounts to the number of extra ChIP-seq samples gained by sharing information between genomic intervals, and the use of these "samples" typically contributes significantly to the estimation of variances for individual intervals. Note also that MAnorm2 empirically determines $d_0$ based on the observed data, which renders the method adaptive to the regularity of variance structure associated with the specific data set (see the section below and also Supplementary Note 2).

For the following differential tests, we further assume that the unscaled variances of non-differential genomic intervals remain invariant across the two groups of samples. Formally, for each interval $i$ that satisfies $\mu_{i,1} = \mu_{i,2}$, we assume $\sigma_{i,1}^2$ equals $\sigma_{i,2}^2$ with a probability of one (i.e., they could be treated as the same random variable). This assumption is consistent with the fact that $\sigma_{i,1}^2$ and $\sigma_{i,2}^2$ follow the same prior distribution as long as $\mu_{i,1} = \mu_{i,2}$. Finally, MAnorm2 tests the null hypothesis $H_0 : \mu_{i,1} = \mu_{i,2}$ for each interval $i$ by using the following key statistic:

$$\tilde{T}_i = \frac{\widehat{\mu_{i,2}} - \widehat{\mu_{i,1}}}{\sqrt{\left(\frac{\gamma_1}{1^T S_{i,1}^{-1} 1} + \frac{\gamma_2}{1^T S_{i,2}^{-1} 1}\right)\tilde{\sigma}_i^2}} \tag{4}$$

where

$$\tilde{\sigma}_i^2 = \frac{d_0 f\left(\frac{\widehat{\mu_{i,1}} + \widehat{\mu_{i,2}}}{2}\right) + (m_1 - 1)\widehat{\sigma_{i,1}^2} + (m_2 - 1)\widehat{\sigma_{i,2}^2}}{d_0 + m_1 + m_2 - 2} \tag{5}$$

According to the theoretical deduction presented in Smyth et al. [1], we assert that $\tilde{T}_i$, termed moderated *t*-statistic, follows a *t*-distribution under the null hypothesis with $(d_0 + m_1 + m_2 - 2)$ degrees of freedom, disregarding the uncertainty associated with the mean estimate (i.e., $\left(\widehat{\mu_{i,1}} + \widehat{\mu_{i,2}}\right)/2$ ). Many existing methods derive the mean signal intensities for determining prior variances (or dispersions, if the negative binomial distribution is used) by taking the average signals across all individual samples [4, 5], which could lead to unbalanced statistical power for identifying up-regulated signals for the two conditions, especially when the numbers of samples belonging to the two conditions differ dramatically from each other. To alleviate this effect, MAnorm2 chooses to take the average signal intensities across conditions, which is especially helpful for balancing the statistical power when $d_0$ is significantly larger than both $m_1$ and $m_2$.

The resulting moderated *t*-statistics could be used to effectively rank genomic intervals in order of statistical evidence of having differential signals between the two conditions. MAnorm2 also gives the exact (two-sided) *p*-values by

$$p_i = 2 \cdot T_{d_0 + m_1 + m_2 - 2}\left(-\left|\tilde{T}_i\right|\right) \tag{6}$$

where $T_N(\cdot)$ refers to the cumulative distribution function of the *t*-distribution with $N$ degrees of freedom.

#### 1.2 Mean-variance curve fitting and parameter estimation

This section discusses the estimation of $f$, $d_0$, $\gamma_1$ and $\gamma_2$ in detail.

To fit the underlying MVC in an unbiased manner, MAnorm2 calculates mean and variance estimates separately within each group of samples and pools the resulting mean-variance pairs into a regression procedure, which is different from many previous methods that use the average signal intensities across all the samples under analysis to fit a curve [3-5]. To make the variance estimates comparable between the two groups of samples being compared, we first deduce an estimate of $\gamma_2/\gamma_1$. Given the model formulation presented in the previous section, we have $\frac{\widehat{t_{i,2}}/\gamma_2}{\widehat{t_{i,1}}/\gamma_1} \sim F_{m_2-1,m_1-1}$ for each genomic interval $i$ that satisfies $\mu_{i,1} = \mu_{i,2}$, where $F_{N_1,N_2}$ refers to the *F*-distribution with $N_1$ and $N_2$ degrees of freedom. And we give an estimator of $\gamma_2/\gamma_1$ as

$$\widehat{\gamma_2/\gamma_1} = \frac{median_i\left(\widehat{t_{i,2}}/\widehat{t_{i,1}}\right)}{F_{m_2-1,m_1-1}^{-1}\left(1/2\right)} \tag{7}$$

where $F_{N_1,N_2}^{-1}(\cdot)$ denotes the inverse of the cumulative distribution function of the corresponding *F*-distribution. Here we use median instead of mean to perform the estimation because the former is more robust to the existence of differential intervals. To further improve the unbiasedness of the estimator, we calculate the median by using only the genomic intervals that are occupied by both groups of samples (see Methods in the main text for a detailed explanation of occupancy states of genomic intervals).

MAnorm2 next pools the mean-variance pairs of the form $\left(\widehat{\mu_{i,1}}, \widehat{t_{i,1}}\right)$ or $\left(\widehat{\mu_{i,2}}, \frac{\widehat{t_{i,2}}}{\widehat{\gamma_2/\gamma_1}}\right)$ into a weighted gamma-family regression procedure, with $(m_1 - 1)$ and $(m_2 - 1)$ as the weights of observations from group $1$ and $2$, respectively. Note that, to enhance the regularity of the set of observations on which the regression is performed, MAnorm2 selects for each group of samples only the genomic intervals occupied by it to calculate mean-variance pairs. Currently, we have devised two candidate schemes for performing the regression. One of them uses a theoretically derived parametric form to fit a generalized linear model (see the following section for the specific form as well as its deduction). This method is most suited to data sets with a highly regular variance structure, where the mean-variance relationship could be expected to be well profiled by the presumed formula (e.g., when the samples of each group are biological replicates of the same experiment). The other method for performing the regression adopts a local regression procedure implemented in the *locfit* package [6]. This method allows for more general mean-variance relationships and is more adaptive to the specific data set [7]. In the study, the parametric method was applied when the ChIP-seq samples grouped together were biological replicates of the same experiment, and we used local regression otherwise. Whichever method is chosen, MAnorm2 iteratively fits an MVC and detects outliers by using 1e-4 and 15 as the lower and upper bounds of residuals (i.e., the ratios of observed variances to fitted ones), respectively [5]. Outliers detected in a round of iteration are removed from the fitting of MVC in the next round of iteration, and the whole regression process finishes as soon as the set of outliers fixes.

In the rest of this section, we ignore the uncertainty associated with the estimate of $f$ , since it is typically fitted on a great number of observations. The method for estimating $d_0$ is similar to the one described in Smyth et al. [1], except that MAnorm2 integrates two estimation results derived from the two groups of samples being compared, respectively. Formally, we define $z_{i,j} = \log \frac{\widehat{t_{i,j}}}{f\left(\widehat{\mu_{i,j}}\right)}$ . Given $\frac{\widehat{t_{i,j}}}{\gamma_j f\left(\mu_{i,j}\right)} \sim F_{m_j-1,d_0}$ , which could be deduced from equation (2) and (3), we assert that the marginal distribution of $z_{i,j}$ is a scaled Fisher's *z*-distribution plus a constant [8], disregarding the uncertainty associated with $\widehat{\mu_{i,j}}$ . And we have

$$\begin{aligned} E\left[z_{i,j}\right] &\approx \log\gamma_j + \psi\left(\frac{m_j-1}{2}\right) - \psi\left(\frac{d_0}{2}\right) + \log\frac{d_0}{m_j-1} \\ \operatorname{var}\left[z_{i,j}\right] &\approx \psi'\left(\frac{m_j-1}{2}\right) + \psi'\left(\frac{d_0}{2}\right) \end{aligned} \tag{8}$$

where $\psi(\cdot)$ and $\psi'(\cdot)$ are the digamma and trigamma functions, respectively.

MAnorm2 next uses these two moments to estimate $\gamma_j$ and $d_0$, respectively.

Noticing that the $z_{i,j}$ from the same group of samples (approximately) have the same expectation and variance values, we give, using the data associated with each group $j$, an estimate of $\psi'(d_0/2)$ by

$$D_j = \frac{\sum_i \left( z_{i,j} - \sum_{i'} z_{i',j} / n_j \right)^2}{n_j - 1} - \psi'\left( \frac{m_j - 1}{2} \right) \tag{9}$$

Note that MAnorm2 only uses the genomic intervals that have been used for fitting $f$ to estimate $\gamma_j$ and $d_0$. Therefore, each of the sum operators in equation (9) is applied only to the intervals whose mean-variance pairs in group $j$ have been involved in the regression process, and $n_j$ denotes the number of such intervals. $n_j$ could vary with $j$, as MAnorm2 selects only the occupied genomic intervals for each group of samples to fit $f$. The final estimate of $d_0$ is obtained by solving

$$\psi'\left( \frac{\widehat{d_0}}{2} \right) = \frac{\sum_j \left( n_j - 1 \right) D_j}{\sum_j \left( n_j - 1 \right)} \tag{10}$$

whose right-hand side has a form similar to the pooled variance estimate in a two-sample *t*-test. Note that $\widehat{d_0}$ is set to positive infinity if the right-hand side of equation (10) is less than or equal to 0, since in this case there is no evidence supporting the variation of the underlying variance (i.e., $\sigma_{i,j}^2$) across the genomic intervals that have the same mean signal intensity. Note also that the marginal distribution of each $\widehat{t_{i,j}}$ is a scaled chi-squared distribution when $d_0$ is positive infinity (as the number of denominator degrees of freedom of an *F*-distribution approaches positive infinity, the *F*-distribution converges to a scaled chi-squared distribution). This fact is consistent with the use of a gamma-family regression procedure to fit $f$.

Finally, we derive for each group $j$ an estimate of $\gamma_j$ by

$$\widehat{\gamma_j} = \exp\left\{\frac{\sum_i z_{i,j}}{n_j} - \psi\left(\frac{m_j - 1}{2}\right) + \psi\left(\frac{\widehat{d_0}}{2}\right) - \log\frac{\widehat{d_0}}{m_j - 1}\right\} \tag{11}$$

Again, the above sum operator is applied only to the genomic intervals whose mean-variance pairs in group $j$ have been used for fitting $f$.

**1.3 Deducing a parametric form for the mean-variance curve**

MAnorm2 deduces an explicit formula that profiles the mean-variance trend by assuming a quadratic relationship between the expectation and the variance of a read count variable, which has been proposed by several previous studies [3, 5, 9]. Formally, suppose $Y$ is a random variable standing for a read count and that it satisfies

$$\mathrm{var}[Y] = \beta_0 E^2[Y] + \beta_1 E[Y] \tag{12}$$

Of note, the whole framework designed in MAnorm2 is for analyzing continuous measurements, which are typically obtained by applying log2 transformation to read counts. We next deduce an approximate formula that connects the variance of $\log_2 Y$ with its expectation by using the delta method [10]. Formally, by defining $X = \log_2 Y$ and investigating its one-order Taylor expansion at $Y = E[Y]$, we have $X \approx \log_2 E[Y] + (Y - E[Y]) \cdot \frac{dX}{dY}|_{Y=E[Y]}$. It follows that

$$E[X] \approx \log_2 E[Y] \tag{13}$$

$$\mathrm{var}[X] \approx \frac{\mathrm{var}[Y]}{E^2[Y]} \cdot \frac{1}{(\log 2)^2} \tag{14}$$

Finally, by substituting the right-hand side of equation (12) for $\mathrm{var}[Y]$ in (14) and using (13) to replace $E[Y]$, we have

$$\mathrm{var}[X] \approx \beta_0' + \beta_1' 2^{-E[X]} \tag{15}$$

where $\beta_i' = \frac{\beta_i}{(\log 2)^2}$ for $i = 0,1$. This form shall be used by MAnorm2 for performing a gamma-family generalized linear regression with identity link.

**Supplementary Note 2. Being adaptive to the regularity of variance structure**

MAnorm2 introduces the notion of prior degrees of freedom in the modeling of mean-variance trend, which is for achieving a wide applicability to data sets of various characteristics. Here, by characteristics of a data set we specifically refer to the regularity of the underlying variance structure, which we assess by measuring the variation in variance residual (from the regression of variance on mean signal intensity) across different genomic intervals (i.e., the variation in $z_{i,j}$ across different $i$ for some fixed $j$; see Supplementary Note 1). For clarification, if a data set is associated with a low level of such variation, it is considered to be of high regularity. Note that MAnorm2 typically derives a large number of prior degrees of freedom for the data sets of high regularity (see equation (9) and (10) in Supplementary Note 1).

In practice, regularity of variance structure as well as the number of prior degrees of freedom derived by MAnorm2 (denoted by $\widehat{d_0}$ in the following) varies significantly across different data sets. This is because there are quite a few factors that could influence the variability of ChIP-seq signals across samples [11], such as technical noise, biological variation, batch effects and so on. As more such factors are involved in a data set, it becomes harder to predict the variances for individual genomic intervals depending solely on their mean signal intensities, and the associated $\widehat{d_0}$ decreases accordingly. For example, we have performed all possible pairwise comparisons of H3K4me3 ChIP-seq signals among four lymphoblastoid cell lines (LCLs; Supplementary Fig. 3). In this analysis, each group of ChIP-seq samples belonging to the same biological condition is comprised of biological replicates for an individual LCL, and the associated variance structure could be expected to be highly regular. As a result, $\widehat{d_0}$ derived by MAnorm2 for these comparisons range from 9.5 to 137.5 and have a median of 18.8. For contrast, we have also made a between-sex comparison of H3K4me3 levels by classifying the four LCLs into two males and two females (Supplementary Fig. 1D). Note that, for this analysis, we created a reference H3K4me3 profile for each individual LCL by taking the average ChIP-seq signals across its biological replicates (see Methods in the main text for details about reference profiles). Compared with the previous scenario, in this case the ChIP-seq profiles grouped together are additionally associated with the epigenetic variation across human individuals [12] and, thus, the associated variance structure should become less regular. Consistently, MAnorm2 derived a $\widehat{d_0}$ of 4.2.

MAnorm2 improves its adaptivity to the specific data set under analysis by empirically estimating $d_0$ and using the estimation result to determine the relative contributions of observed variances and prior ones to the final variance estimates for individual genomic intervals (see equation (5) in Supplementary Note 1). Based on the corresponding gene expression data, we compared the performance of MAnorm2 with two of its variants in the above two scenarios of group-level differential ChIP-seq analysis. Note that we used the comparison between GM12890 and SNYDER LCLs (2 vs. 2 biological replicates) as representative of the first scenario, as the comparison is associated with the same group sizes as the between-sex comparison (2 vs. 2 LCLs). The two variants of MAnorm2, referred to as no-MVC and MVC-only, utilize different strategies from MAnorm2 to derive final variance estimates for individual intervals. More specifically, no-MVC and MVC-only directly use observed and prior variances as the final variance estimates, respectively, while MAnorm2 integrates the two types of variances by taking a weighted average, with the weights depending on the $d_0$ estimated from the data (technically, no-MVC and MVC-only are equivalent to always treating $d_0$ as 0 and positive infinity, respectively). As described above, the underlying variance structure in the first scenario is of high regularity, and there is a low variability in variance across the genomic intervals having the same mean signal intensity (Supplementary Fig. 1A). In this case, prior variances alone could serve as reliable variance estimates for individual intervals. In contrast to the prior variances, the observed variances in this scenario are associated with large uncertainty due to the small numbers of biological replicates, which results in a significantly worse performance of no-MVC compared with MVC-only (Supplementary Fig. 1C). As for MAnorm2, it derived a $\widehat{d_0}$ of 14.6, which is more than seven times the number of observed degrees of freedom (i.e., the number of free measurements of signal intensities for calculating the observed variance for each genomic interval, which equals the total number of ChIP-seq samples minus the number of groups of samples). Consequently, the final variance estimates used by MAnorm2 are dominated by the prior variances, and MAnorm2 therefore exhibits virtually the same performance as MVC-only. In comparison, the variance structure in the second scenario is considerably less regular, and MAnorm2 accordingly derived a $\widehat{d_0}$ of 4.2 (Supplementary Fig. 1B), which is comparable to the number of observed degrees of freedom. In this scenario, no-MVC continues to suffer from small group sizes, and MVC-only completely ignores the high fluctuation of the variances of the genomic intervals having the same mean signal intensity. As a result, both no-MVC and MVC-only are clearly outperformed by

MAnorm2 (Supplementary Fig. 1D). Together, these results demonstrate the adaptivity of MAnorm2 and suggest a wide applicability of it.

**Supplementary Note 3. Simultaneously comparing multiple groups of ChIP-seq samples**

This note gives a formal description of the statistical model designed in MAnorm2 for simultaneously comparing more than two groups of ChIP-seq samples corresponding to different biological conditions. We keep using the notations defined in Supplementary Note 1, except that the group index $j$ now takes integers from 1 through $C$, where $C$ is the total number of groups of samples to be compared. To be rigorous, we give a succinct but still self-contained description of the associated model formulation and hypothesis testing.

For each genomic interval $i$ in each group $j$, we assume

$$X_{i,j} \Big| \sigma_{i,j}^2 \sim MVN\left(1\mu_{i,j}, S_{i,j}\left(\gamma_j \sigma_{i,j}^2\right)\right)$$
$$\frac{1}{\sigma_{i,j}^2} \sim \frac{1}{f\left(\mu_{i,j}\right)} \cdot \frac{\chi_{d_0}^2}{d_0} \tag{16}$$

We further assume that the unscaled variance of each non-differential genomic interval remains invariant across groups. Formally, for each interval $i$ that satisfies $\mu_{i,1} = \mu_{i,2} = \ldots = \mu_{i,C}$, we assume $\sigma_{i,1}^2 = \sigma_{i,2}^2 = \ldots = \sigma_{i,C}^2$ happens with a probability of one (i.e., they could be treated as the same random variable). This assumption is consistent with the fact that $\sigma_{i,1}^2, \sigma_{i,2}^2, \ldots, \sigma_{i,C}^2$ follow the same prior distribution as long as $\mu_{i,1} = \mu_{i,2} = \ldots = \mu_{i,C}$. For later use, we derive expressions of mean and variance estimators by applying the generalized least squares method:

$$\widehat{\mu_{i,j}} = \left(1^T S_{i,j}^{-1} 1\right)^{-1} 1^T S_{i,j}^{-1} X_{i,j}$$
$$\widehat{t_{i,j}} = \frac{\left(X_{i,j} - 1\widehat{\mu_{i,j}}\right)^T S_{i,j}^{-1} \left(X_{i,j} - 1\widehat{\mu_{i,j}}\right)}{m_j - 1} \tag{17}$$

Methods detailed in Supplementary Note 1 for estimating $f$, $d_0$ and $\gamma_j$ could be naturally extended to comparisons of more than two groups of samples with few modifications. Specifically, for fitting $f$, we select one from the $C$ groups as baseline and derive an estimate of $\gamma_j / \gamma_b$ for each $j \neq b$ by using equation (7), where $b$ refers to the selected baseline group. Then, the mean-variance pairs having a form of $\left(\widehat{\mu_{i,b}}, \widehat{t_{i,b}}\right)$ or $\left(\widehat{\mu_{i,j}}, \frac{\widehat{t_{i,j}}}{\widehat{\gamma_j / \gamma_b}}\right)$ for $j \neq b$ are pooled into a weighted gamma-family regression procedure, with $\left(m_j - 1\right)$ as the weight of observations from group $j$. For the selection of baseline group, MAnorm2 utilizes an algorithm similar to the one for selecting a baseline ChIP-seq sample for normalizing a group of samples (see Methods in the main text). More specifically, it picks out the genomic intervals that are occupied by all the $C$ groups of samples and uses the $\widehat{t_{i,j}}$ of them to construct a matrix, whose rows and columns correspond to the intervals and the groups, respectively. MAnorm2 then applies the median-ratio strategy [7] to the matrix and derives the “size factor” of each group. Finally, the group whose log2 size factor is closest to 0 is selected as baseline. After fitting $f$, the estimation of $d_0$ and $\gamma_j$ is accomplished by using equation (9), (10) and (11). The resulting estimates of $f$, $d_0$ and $\gamma_j$ are treated as non-stochastic in the subsequent statistical tests, as they are typically derived based on a great number of observations.

We next detail the procedure for testing the null hypothesis $H_0 : \mu_{i,1} = \mu_{i,2} = \ldots = \mu_{i,C}$ for each genomic interval $i$. As in the one-way analysis of variance (ANOVA), we first fit the full model and calculate the corresponding residual sum of squares (RSS):

$$RSS_i = \sum_{j=1}^{C} \frac{\left(m_j - 1\right)\widehat{t_{i,j}}}{\gamma_j} \tag{18}$$

We then fit a reduced model by assuming all the $C$ biological conditions are associated with the same mean signal intensity in interval $i$:

$$\mu_i^{(0)} = \frac{\sum_{j=1}^{C} \left(\frac{1^T S_{i,j}^{-1} 1}{\gamma_j}\right) \widehat{\mu_{i,j}}}{\sum_{j=1}^{C} \frac{1^T S_{i,j}^{-1} 1}{\gamma_j}} \tag{19}$$

where $\mu_i^{(0)}$ is intrinsically a weighted average of mean estimates across conditions, with the weights being inversely proportional to their variances. And the associated RSS could be derived by

$$RSS_i^{(0)} = \sum_{j=1}^{C} \frac{\left(X_{i,j} - 1\mu_i^{(0)}\right)^T S_{i,j}^{-1} \left(X_{i,j} - 1\mu_i^{(0)}\right)}{\gamma_j} \tag{20}$$

Before defining the final key statistic for testing the null hypothesis for interval $i$, we summarize some facts regarding the distributions of associated random variables as follows. Under the $H_0$, we have $\mu_{i,1} = \mu_{i,2} = \ldots = \mu_{i,C}$ (denoted by $\mu_i$ in the following) and that $\sigma_{i,1}^2, \sigma_{i,2}^2, \ldots, \sigma_{i,C}^2$ refer to the same random variable (denoted by $\sigma_i^2$ in the following). Based on equation (16) and previous studies of one-way ANOVA, we have (under the $H_0$)

$$\begin{aligned} &\frac{1}{\sigma_i^2} \sim \frac{1}{f(\mu_i)} \cdot \frac{\chi_{d_0}^2}{d_0} \\ &RSS_i \middle| \sigma_i^2 \perp \left(RSS_i^{(0)} - RSS_i\right) \middle| \sigma_i^2 \\ &RSS_i \middle| \sigma_i^2 \sim \sigma_i^2 \cdot \chi_{\sum_j m_j - C}^2 \\ &\left(RSS_i^{(0)} - RSS_i\right) \middle| \sigma_i^2 \sim \sigma_i^2 \cdot \chi_{C-1}^2 \end{aligned} \tag{21}$$

in which the second formula suggests that the two random variables are conditionally (on $\sigma_i^2$) independent of each other. Equation (21) gives all the results that are necessary for deriving

$$\frac{\left(RSS_i^{(0)} - RSS_i\right) \Big/ (C-1)}{\left(RSS_i + d_0 f(\mu_i)\right) \Big/ \left(\sum_j m_j - C + d_0\right)} \sim F_{C-1, \sum_j m_j - C + d_0} \tag{22}$$

Finally, we define a moderated *F*-statistic for interval $i$ as

$$\tilde{F}_i = \frac{\left(RSS_i^{(0)} - RSS_i\right) \Big/ (C-1)}{\left(RSS_i + d_0 f\left(\sum_j \widehat{\mu_{i,j}} \big/ C\right)\right) \Big/ \left(\sum_j m_j - C + d_0\right)} \tag{23}$$

which approximately follows $F_{C-1, \sum_j m_j - C + d_0}$ under the null hypothesis, considering the uncertainty of the mean estimate for deducing the prior variance of interval $i$ (i.e., $\sum_j \widehat{\mu_{i,j}} \big/ C$). $\tilde{F}_i$ has a form similar to the classical *F*-statistic in one-way ANOVA, except that its variance estimate (i.e., the denominator) has incorporated additional information regarding $\sigma_i^2$ , which is exactly obtained by modeling mean-variance dependence. This incorporation of prior variances helps stabilizing the variance estimates for individual intervals as well as increasing the statistical power for identifying differential intervals across conditions, which could be reflected by the increased number of denominator degrees of freedom of $\tilde{F}_i$ compared to the classical *F*-statistic. Note also that $\tilde{F}_i$ is similar to the moderated *F*-statistic designed in Smyth et al. [1], except that the latter uses a constant prior variance for all genomic intervals and does not take mean-variance dependence into account. As explained in Supplementary Note 1, here we derive the mean estimates for determining prior variances by taking the average signal intensities across conditions rather than individual samples, which is for avoiding biasing the mean estimates towards the conditions that are associated with more ChIP-seq samples than the others. In practice, such biases typically lead to stronger statistical power for identifying up-regulated signals in the conditions with more samples. Taking the average signals across conditions is especially effective for alleviating the unbalanced statistical power when $d_0$ is significantly larger than $\left(\sum_j m_j - C\right)$.

Accordingly, MAnorm2 gives the *p*-value of the statistical test for interval $i$ by

$$p_i = 1 - F_{C-1, \sum_j m_j - C + d_0}\left(\tilde{F}_i\right) \tag{24}$$

where $F_{N_1,N_2}\left(\cdot\right)$ refers to the cumulative distribution function of the *F*-distribution with $N_1$ and $N_2$ degrees of freedom.

### Supplementary Figure Legends

**Supplementary Figure 1. Being adaptive to the complexity of variance structure across different scenarios. (A)** Scatter plot showing the mean-variance trend associated with the comparison of H3K4me3 levels between GM12890 and SNYDER lymphoblastoid cell lines (LCLs). Red line depicts the fitted mean-variance curve (MVC), and $d_0$ gives the corresponding number of prior degrees of freedom. **(B)** Scatter plot for the comparison of H3K4me3 levels between male and female LCLs. Note that both comparisons are two-versus-two and, thus, the dispersion level of observed mean-variance pairs around MVC is comparable between the two scenarios. **(C, D)** For each of the two differential analyses, the proportion of true discoveries among top ranked genomic intervals at gene promoters is plotted against the number of top ranked intervals. Here true discoveries are defined as the intervals that are linked with differentially expressed genes (DEGs), which are identified by applying DESeq2 to the corresponding RNA-seq data with a *p*-value cutoff of 0.01.

**Supplementary Figure 2. Comparing MAnorm2 with existing empirical Bayes methods that model mean-variance/dispersion relationship by applying them to comparisons of H3K4me3 ChIP-seq signals between a pair of LCLs.** We have performed all possible pairwise comparisons among GM12890, GM12891, GM12892 and SNYDER LCLs. For each comparison, the proportion of true discoveries among top ranked genomic intervals at gene promoters is plotted against the number of top ranked intervals.

**Supplementary Figure 3. Comparing MAnorm2 with other tools for group-level differential ChIP-seq analysis by applying them to comparisons of H3K4me3 ChIP-seq signals between a pair of LCLs.** We have performed all possible pairwise comparisons among GM12890, GM12891, GM12892 and SNYDER LCLs. For each comparison, the proportion of true discoveries among top ranked genomic intervals at gene promoters is plotted against the number of top ranked intervals.

### Supplementary Figures

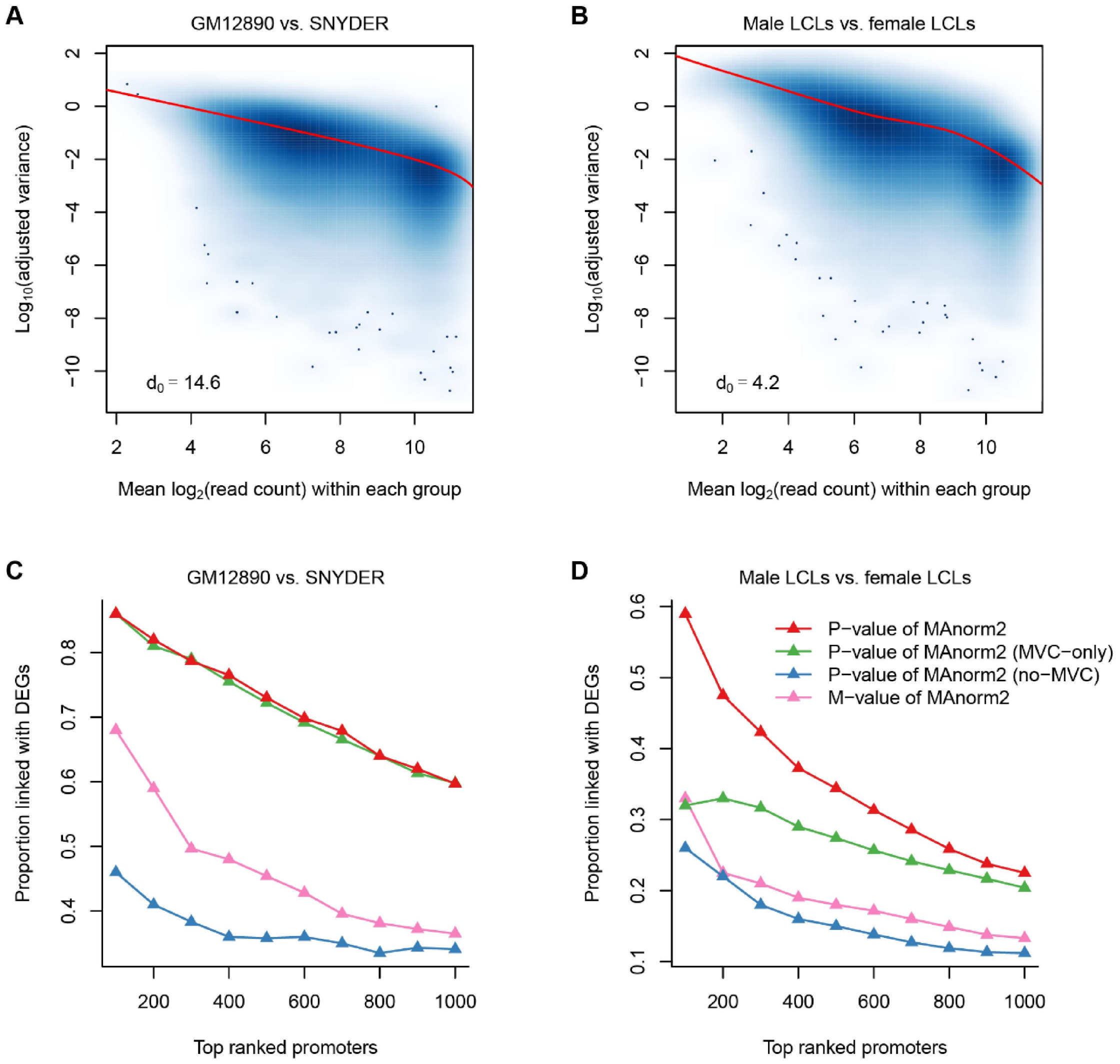


**Supplementary Figure 1**

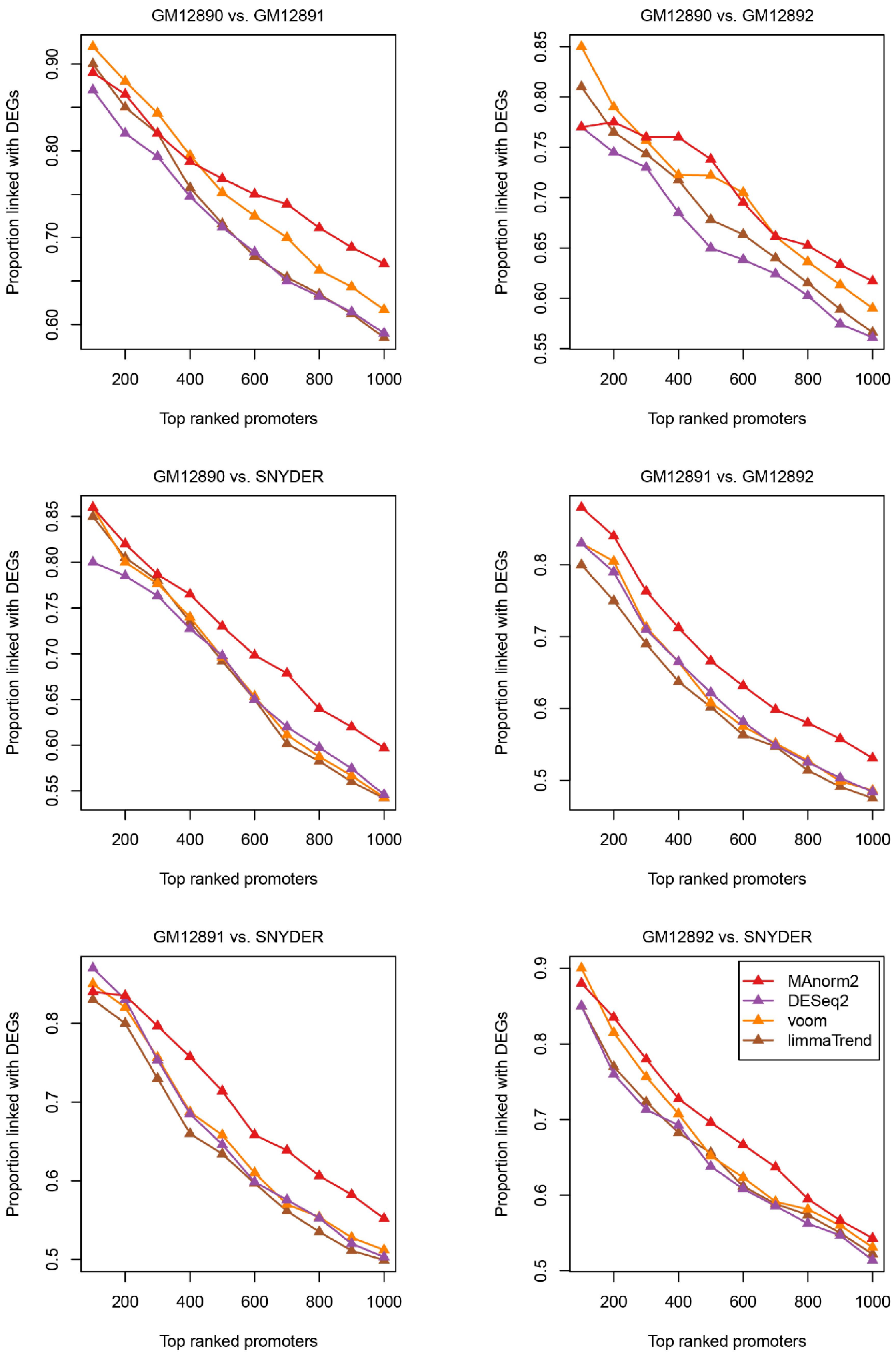


**Supplementary Figure 2**

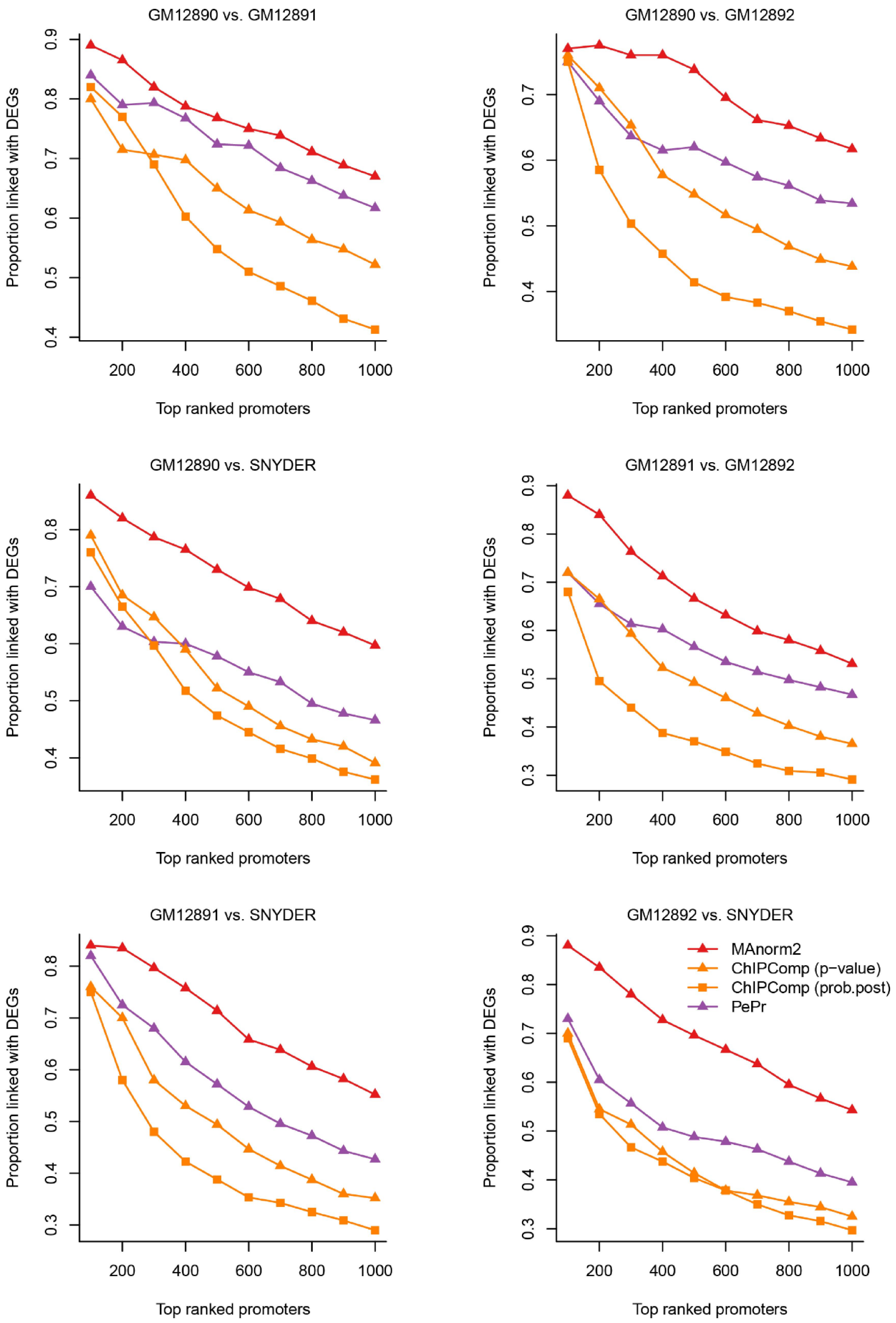


**Supplementary Figure 3**
